# How ascorbate tunes the activity and substrate selectivity of Fe(II)-dependent dioxygenase superfamily enzymes

**DOI:** 10.1101/2025.05.06.652568

**Authors:** Lucas O. Calzini, Marcin Warminski, Joanna Kowalska, Jacek Jemielity, Jeffrey S. Mugridge

**Author notes:** Corresponding Author. Department of Chemistry & Biochemistry, University of Delaware, Newark, DE 19716, United States.

## Abstract

Across all domains of life, Fe(II)- and 2-oxoglutarate(2-OG)-dependent dioxygenase (FODD) superfamily enzymes carry out pivotal oxidation reactions that underlie key biological processes ranging from hormone biosynthesis to oxygen sensing to DNA repair and RNA modification. This study combines enzymology and structural biology to elucidate a new mechanism of FODD regulation whereby cofactor ascorbate (vitamin C) concentrations tune both the activity and substrate selectivity of FODD enzymes involved in RNA demethylation, and for the first time reveals the structural basis for ascorbate’s interaction with the FODD superfamily active site. Because ascorbate concentrations vary by over 100-fold across different cell types and disease states, our mechanistic work demonstrates how ascorbate levels likely play a critical, but underappreciated role in regulating RNA modification across the epitranscriptome and, more broadly, in regulating diverse biological oxidation reactions across the cell and human diseases.

## INTRODUCTION

The **<u>F</u>**e(II)- and 2-**<u>o</u>**xoglutarate-**<u>d</u>**ependent **<u>d</u>**ioxygenase (FODD) enzyme superfamily (**Figure 1a**) catalyzes essential oxidation reactions on a wide range of peptide and nucleic acid substrates throughout the cell.^1–4^ FODDs play key roles in collagen formation,^5,6^ fatty acid metabolism,^7,8^ oxygen sensing and hypoxia response,^9–11^ DNA damage repair,^12–14^ and both epigenetic (DNA)^15–19^ and epitranscriptomic (RNA) modifications.^20–22^ Dysregulation of FODDs has been directly linked to the development and progression of diverse human cancers^23–26^ – including leukemia,^27,28^ gliomas,^29–31^ carcinoma,^32^ lung,^33^ and breast^34–36^ cancers – as well as neurodegenerative diseases^37,38^ and metabolic disorders.^39–41^

**Figure 1.**
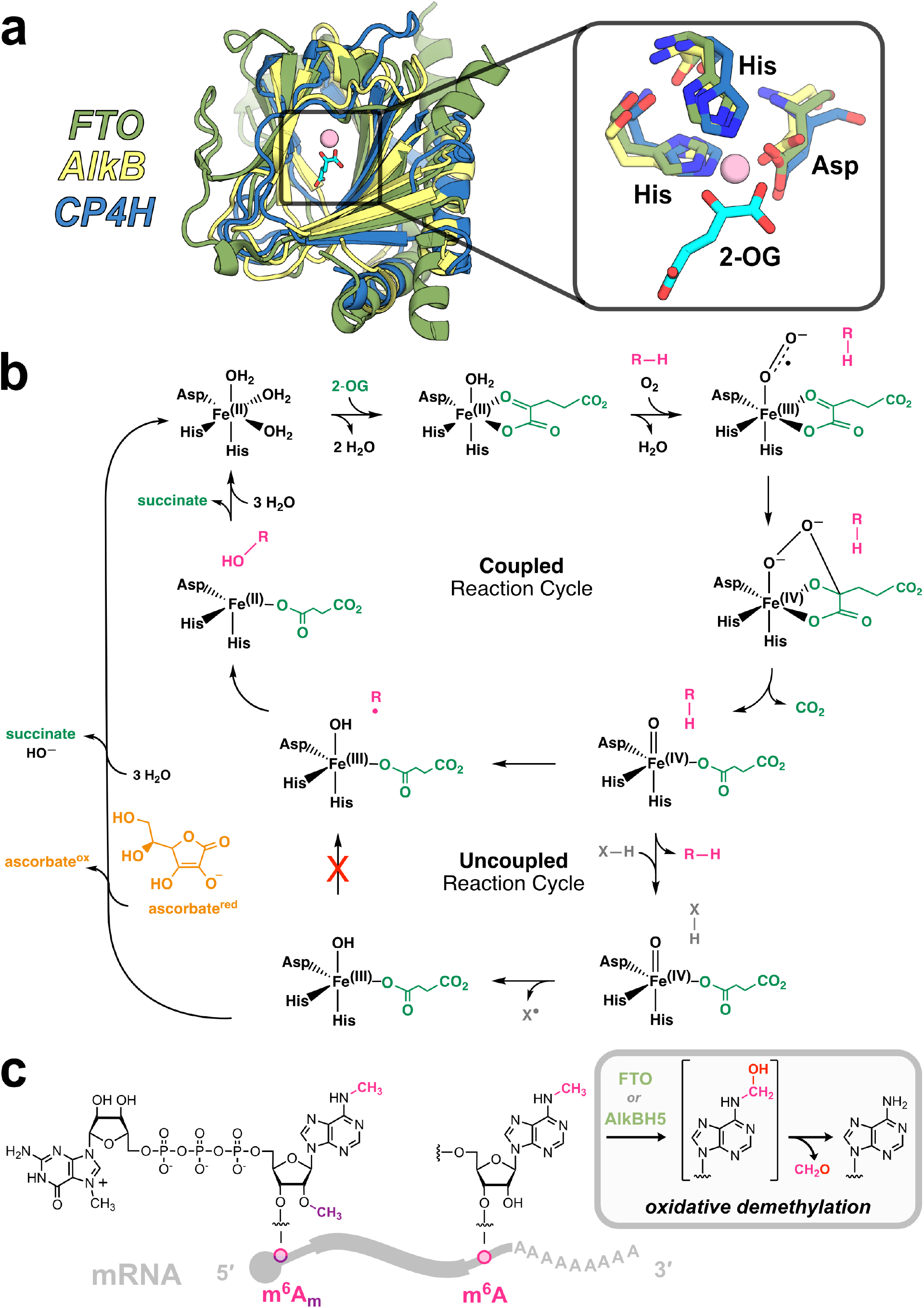
FODD structure, catalytic cycle, and RNA demethylation reactions. **(a)** Structural alignment of FODD catalytic domains of FTO (PDB 4ZS2), AlkB (PDB 3RZJ) and CP4H (AlphaFold 3 prediction^66^), with inset showing conserved His-Asp-His active site residues that coordinate the Fe(II) metal center along with cofactor 2-OG. Although these FODDs oxidize very different substrates (RNA for FTO, DNA for AlkB, peptide for CP4H), their catalytic domains have a high degree of structural homology. **(b)** Canonical, proposed catalytic cycle for FODDs. The coupled reaction cycle begins with the binding of 2-OG to the Fe(II) center, followed by oxygen activation, and then 2-OG decarboxylation, producing CO_2_ and succinate. This decarboxylation reaction (green) also generates the highly reactive Fe(IV)-oxo, which hydroxylates the prime substrate (R–H) through a radical rebound mechanism (pink). The uncoupled reaction cycle occurs when 2-OG decarboxylation and Fe(IV)-oxo generation is not coupled to prime substrate hydroxylation. In this case, the reactive Fe(IV)-oxo abstracts a hydrogen from its environment and eventually stalls in the Fe(III)-hydroxyl state. Ascorbate is then thought to act as a reducing agent to rescue this stalled state and regenerate Fe(II) for further turnovers; however the nature of the Fe-ascorbate interaction, and when and how ascorbate accesses the active site, have remained unclear. **(c)** The *N*^*6*^-methyl groups on abundant mRNA modifications m^6^A_m_ and m^6^A can undergo oxidative demethylation mediated by FODD enzymes FTO and AlkBH5; FTO (acts on m^6^A or m^6^A_m_) and AlkBH5 (acts on m^6^A) first hydroxylate the *N*^6^-methyl group, which then decomposes to adenosine (inset).

The canonical FODD catalytic cycle consists of two coupled chemical reactions^42–46^ (**Figure 1b, ‘coupled reaction cycle’**): (1) the decarboxylation of 2-oxoglutarate (2-OG) to produce succinate (**Figure 1b, green**) and a reactive Fe(IV)-oxo intermediate,^47,48^ followed by (2) the oxidation of prime substrate to form a hydroxylated product via a radical rebound mechanism (**Figure 1b, pink**). However, FODDs can also undergo 2-OG decarboxylation and Fe(IV)-oxo formation that is *not coupled* to prime substrate oxidation, which is thought to generate unreactive Fe(III)-hydroxyl species in the active site, stalling enzyme activity and turnover (**Figure 1b**, ‘uncoupled reaction cycle’). Here, vitamin C (ascorbate) is thought to act as a critical cofactor that reduces these unreactive Fe(III) states back to Fe(II) to rescue enzyme activity.^48–52^ The most well-studied and well-known example of this is collagen prolyl 4-hydroxylase (CP4H), an FODD that hydroxylates proline in collagen precursor peptides and is completely dependent on ascorbate for proper activity.^53^ This example is found in many introductory biochemistry textbooks^54^, because the strict ascorbate dependence of CP4H is a primary reason that people with serious dietary ascorbate deficiencies develop scurvy – loss of CP4H activity results in defects in collagen synthesis.^55^ Limited studies also suggest other FODD enzymes – like the DNA-modifying enzymes TET^56^ and AlkB^57^, as well as RNA demethylase FTO^58^ – have some ascorbate dependence, so researchers have typically carried out all *in vitro* assays with FODDs in the presence of mM ascorbate concentrations to ensure robust activity.

However, we lack a systematic understanding of how ascorbate impacts dioxygenase activity levels across the FODD superfamily, and whether ascorbate requirements vary between different FODD enzymes and their diverse substrates. These are critical questions for FODD regulation, because ascorbate concentrations can be strikingly different between human cell types and disease states – ranging from 0.2 to 20 mM across different human cells and tissues,^52^ with cancerous cells having significantly dysregulated ascorbate levels^11,59,60^ – and these environments may differentially control FODD activity or selectivity. Furthermore, despite the link between ascorbate and FODD activity being known for over 50 years, there is to date almost no biophysical or structural information about how ascorbate specifically engages the FODD active site or what structural features position ascorbate to selectively rescue enzyme activity.

Here, we use RNA demethylases FTO and AlkBH5 as model FODD enzymes to uncover significant differences in the regulation of FODD activity and selectivity by ascorbate. FTO and AlkBH5 are RNA methyl ‘erasers’ that selectively remove the highly abundant *N*^6^-methyladenosine (m^6^A; FTO or AlkBH5) modification found in the body of mRNA and the *N*^*6*^-2′-*O*-dimethyladenosine (m^6^A_m_; FTO only) modification found at the 5′ cap of mRNA (**Figure 1c**).^61–65^ Using enzymology, we show that FTO and AlkBH5 differentially utilize ascorbate during their oxidative demethylation reactions, and that FTO’s ascorbate usage varies dramatically for RNA substrates with different *k*_cat_ values. Further kinetic experiments reveal that FTO, unlike AlkBH5, has a high propensity to undergo 2-OG decarboxylation in the absence of prime substrate, revealing a key factor in understanding its ascorbate dependence. We also present the first structural and biophysical data showing how the FTO active site selectively binds ascorbate to reduce and rescue stalled enzyme states using conserved residues. Together, our results suggest a new model for FODD ascorbate usage and dependence that is determined primarily by: (a) an FODD’s propensity to undergo 2-OG decarboxylation in the absence of prime substrate and (b) the relative efficiency of the catalytic step during coupled reaction cycles. The discovery that RNA demethylases can have dramatically different ascorbate dependencies suggests that cellular ascorbate levels may selectively tune mRNA demethylation across the transcriptome. More broadly, we propose that these principles will be true for many FODD enzymes and that ascorbate levels, which vary across cell types and disease states, could play a key role in selectively regulating the activity and substrate specificity of all FODD enzymes.

## RESULTS

### FODD enzymes FTO and AlkBH5 differentially utilize ascorbate

To investigate ascorbate usage within the Fe(II)- and 2-OG-dependent dioxygenase (FODD) superfamily, we used RNA demethylases FTO and AlkBH5 as model FODD enzymes, since these allow for direct comparison of ascorbate-dependent demethylation activity on the same m^6^A-containing RNA substrates. We performed *in vitro* demethylation assays with FTO and AlkBH5 on either a linear 9-mer m^6^A-RNA or a structured, stem-loop 25-mer m^6^A-RNA substrate, in both the presence or absence of ascorbate (**Figure 2**). Each enzyme (FTO or AlkBH5) was incubated with cofactors Fe(II) and 2-OG along with m^6^A-RNA substrate and either 0 mM or 2 mM ascorbate (**Figure 2a**). After addition of m^6^A-RNA substrate, demethylation kinetics were monitored over time by digesting quenched reaction mixtures to single RNA nucleosides and quantifying the relative abundances of m^6^A and A via UHPLC-MS (**Supplementary Figure 1**). We measured initial demethylation rates over a range of substrate concentrations to determine kinetic parameters for FTO- and AlkBH5-mediated m^6^A demethylation with or without ascorbate (**Supplementary Table 1**). The resulting Michaelis-Menten plots show that FTO exhibits a near complete dependence on ascorbate for efficient demethylation of either the linear 9-mer or structured 25-mer m^6^A-RNA substrates (**Figure 2b,c**). In contrast, AlkBH5 shows nearly the same level of demethylation activity on m^6^A-RNAs in either the presence or absence of ascorbate (**Figure 2d,e**). These kinetic data show that for both structured and unstructured model m^6^A-containing RNA substrates, FODD family demethylases FTO and AlkBH5 have dramatically different ascorbate dependencies – with FTO-mediated demethylation being almost entirely dependent on the presence of cofactor ascorbate, while AlkBH5-mediated demethylation is almost entirely independent of ascorbate.

**Figure 2.**
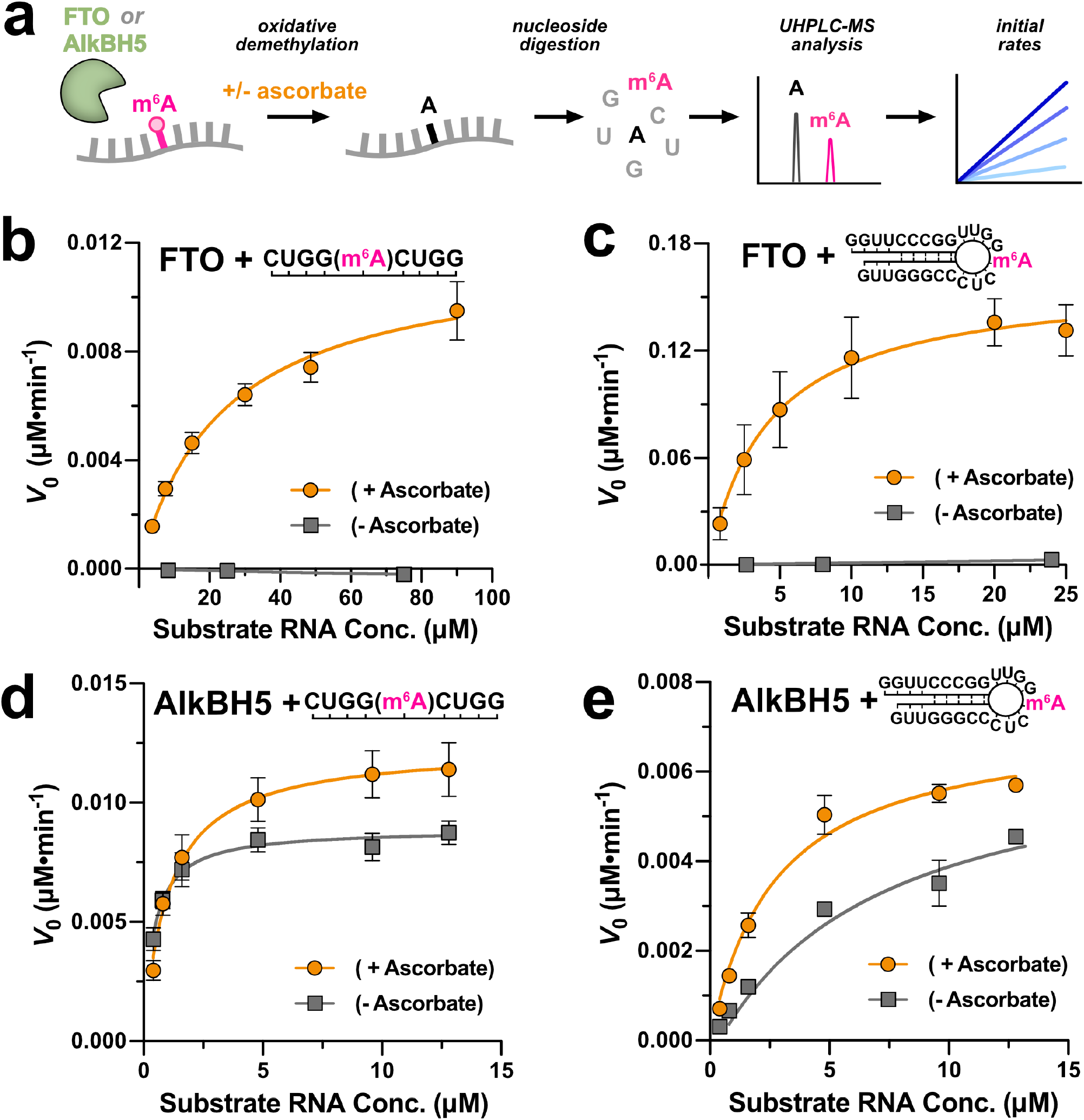
FTO and AlkBH5 have different ascorbate dependencies for their m^6^A demethylation reactions. **(a)** Schematic overview of kinetic assays showing demethylation of an m^6^A-RNA substrate, followed by nucleoside digestion, quantification by UHPLC-MS, and initial rate determination. **(b, c)** Michaelis-Menten plots of initial rate (*V*_0_) versus substrate concentration for *in vitro* FTO-mediated m^6^A-RNA demethylation reactions in the presence (orange) or absence (gray) of ascorbate. FTO (500 nM) was incubated with Fe(II), 2-OG, and varied concentrations of a linear 9-mer m^6^A-RNA (**b**) or a 25-mer stem-loop m^6^A-RNA (**c**) substrate, with either 0 mM (gray) or 2 mM (orange) added ascorbate. **(d, e)** Michaelis-Menten plots of initial rate (*V*_0_) versus substrate concentration for *in vitro* AlkBH5-mediated m^6^A-RNA demethylation reactions in the presence (orange) or absence (gray) of ascorbate. Reactions were carried out similarly to FTO as above, but with 200 nM AlkBH5 for linear 9-mer m^6^A-RNA reactions **(d)** and 100 nM AlkBH5 for 25-mer stem-loop m^6^A-RNA reactions **(e)**. All error bars represent SEM of three replicates. *k*_cat_ and *K*_M_ values determined in these analyses are listed in **Supplementary Table 1**.

### FTO activity is less dependent on ascorbate for efficiently demethylated RNA substrates

We next examined how FTO’s ascorbate dependence varies for demethylation of different RNA substrates. We performed end-point demethylation assays with FTO and linear 9-mer m^6^A-RNA, 25-mer stem-loop m^6^A-RNA, and 5′-cap m^6^A_m_-RNA substrates over a range of ascorbate concentrations to determine FTO’s ascorbate requirements with each substrate (**Figure 3a**). Consistent with our previous experiments, the linear m^6^A-RNA substrate was the most sensitive to diminishing ascorbate concentrations, followed next by the stem-loop m^6^A-RNA. The m^6^A_m_-RNA substrate was the least sensitive, still showing relatively robust FTO-mediated demethylation activity in the complete absence of ascorbate. To explain these trends, we determined kinetic parameters for each substrate (**Supplementary Table 1, Supplementary Figure 2**) and compared their *k*_*cat*_ and *K*_*M*_ values (**Figure 3b**). These data show that while *K*_*M*_ remains roughly the same across all three RNA substrates, *k*_*cat*_ differs dramatically. Furthermore, the relative trend in *k*_*cat*_ values mirrors the trend in ascorbate sensitivity: the m^6^A_m_ cap substrate has the highest *k*_*cat*_ and is the least dependent on ascorbate, whereas the linear m^6^A substrate has the lowest *k*_*cat*_ and is the most dependent on ascorbate. These data strongly suggest that substrates whose catalysis is highly efficient (*i*.*e*. high *k*_cat_ values) require less ascorbate.

**Figure 3.**
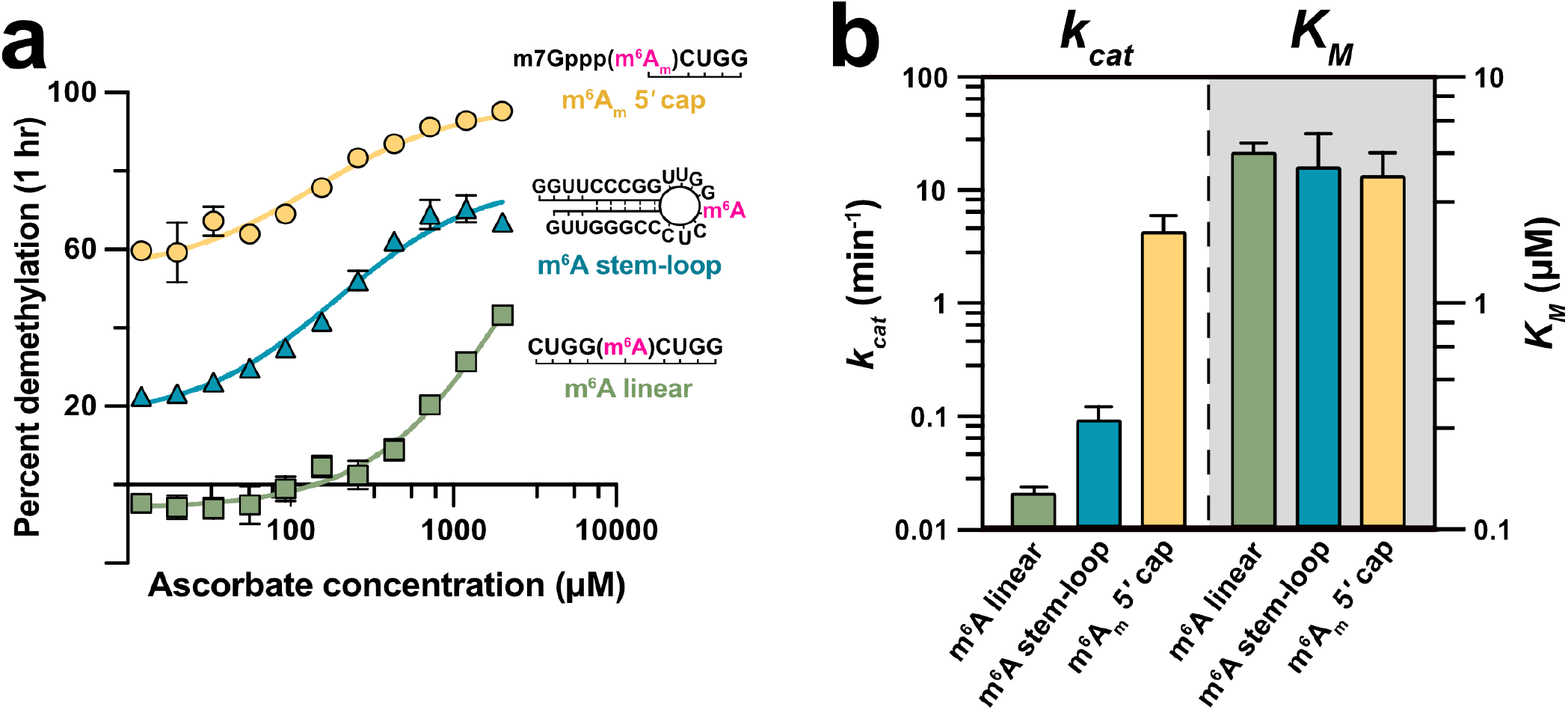
Efficient substrates of FTO are less dependent on ascorbate for demethylation. **(a)** The ascorbate dependence for demethylation of three different FTO substrates – linear 9-mer m^6^A-RNA (green squares), 25-mer stem loop m^6^A-RNA (blue triangles), and 5′-cap m^6^A_m_-RNA (yellow circles) – was evaluated by measuring the percent demethylation of each substrate after 1 hour at varying ascorbate concentrations (0 – 2 mM). The sequence and structure of each RNA oligo substrate is shown. All error bars are the SEM of three replicates. **(b)** Bar graph of kinetic parameters for demethylation of different m^6^A- or m^6^A_m_-RNA substrates by FTO. *k*_*cat*_ values are shown on the left side of the bar graph (white background) and use the left-hand y-axis; *K*_*M*_ values are shown on the right-hand side of the bar graph (gray background) and use the right-hand y-axis. Bar coloring corresponds to the m^6^A- or m^6^A_m_-RNA substrate data shown in **a**. The displayed *k*_*cat*_ and *K*_*M*_ values are derived from fits of initial rate versus substrate concentration data shown in **Figure 2b,c** for linear and stem-loop substrates and **Supplementary Figure 2** for m^6^A_m_ 5′ cap substrate.

### FTO and AlkBH5 have different decarboxylation kinetics in the absence of substrate

Building from our enzymology data above that shows FTO-mediated demethylation is sensitive to ascorbate concentrations, but AlkBH5-mediated reactions are not, we next asked whether these two enzymes have different propensities to undergo uncoupled decarboxylation that might explain their different ascorbate dependencies. To investigate this aspect of the FODD catalytic cycle for FTO and AlkBH5, we monitored the progress of the 2-OG decarboxylation reaction by tracking 2-OG consumption and succinate production over time in the absence of prime substrate via ^1^H-NMR (**Figure 4a**). When FTO was incubated with Fe(II), 2-OG, and ascorbate, we see steady consumption of 2-OG over time and the corresponding production of succinate, indicating that FTO readily catalyzes the decarboxylation reaction in the absence of prime substrate (**Figure 4b,*ii***). This observation is consistent with previous studies of TET1 and other FODDs that can also carry out uncoupled 2-OG decarboxylation.^67,68^ For AlkBH5 on the other hand, very little 2-OG is consumed and very little succinate is produced over the 6-hour time course of the experiment (**Figure 4b,*iii***), indicating that unlike FTO, AlkBH5 cannot readily catalyze the decarboxylation reaction without its prime substrate. Integrating and quantifying the relative 2-OG and succinate changes over time (**Figure 4c,d**) shows that the FTO-catalyzed decarboxylation reaction outpaced that of AlkBH5 by approximately 15-fold. These data reveal FTO’s much stronger propensity for decarboxylation of 2-OG in the absence of prime substrate, in stark contrast to AlkBH5’s relative inability to undergo oxygen activation and 2-OG decarboxylation in the absence of its RNA substrate. These differences in decarboxylation activity suggest a second key predictor of FODD ascorbate dependence: the ability of the enzyme to catalyze decarboxylation in the absence of prime substrate.

**Figure 4.**
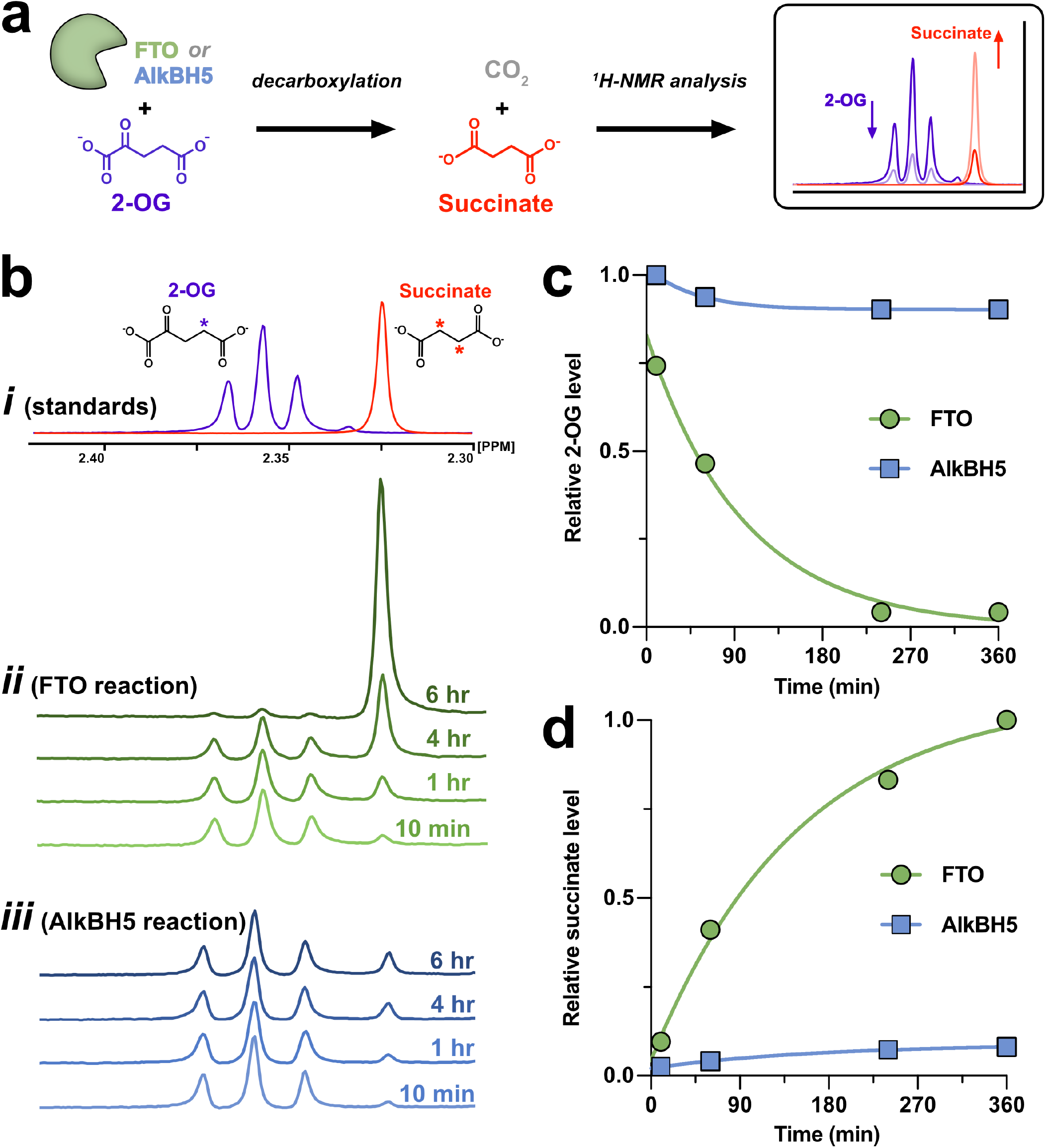
FTO catalyzes 2-OG decarboxylation without prime substrate and AlkBH5 does not. **(a)** Schematic of 2-OG decarboxylation assays, showing how 2-OG and succinate resonances are expected to change during ^1^H-NMR analysis. **(b) *i***, ^1^H-NMR spectra of 2-OG and succinate standards showing resonances used for decarboxylation reaction analysis; resonances correspond to the starred protons on each chemical structure. ***ii***, Decarboxylation reaction of FTO with 2-OG in the absence of prime m^6^A-RNA substrate. The singlet resonance increasing in intensity over time corresponds to the production of succinate, while the triplet resonance decreasing over time corresponds to the consumption of 2-OG. ***iii***, Analogous reaction as in ***ii*** but with AlkBH5, again in the absence of prime m^6^A-RNA substrate. No appreciable decarboxylation activity was measured, as indicated by minimal changes in either the 2-OG or succinate resonances. **(c)** Normalized ^1^H-NMR 2-OG peak areas from reactions in **b *ii*** and ***iii*. (d)** Normalized ^1^H-NMR succinate peak areas from reactions in **b *ii*** and ***iii***.

### Ascorbate selectively stabilizes Fe(II) levels during FTO-mediated demethylation reactions

Previous studies have suggested that ascorbate acts in the FODD catalytic cycle by reducing stalled Fe(III) states back to catalytically-active Fe(II), thus rescuing enzyme that has been shunted into the uncoupled reaction cycle (**Figure 1b**).^48,50,51^ To monitor how ascorbate helps to regulate Fe(II) levels in FTO-mediated catalysis, we ran demethylation reactions with FTO and quenched the reaction at different times with phenanthroline, which strongly chelates iron and selectively absorbs at 511 nm when complexed with Fe(II).^69^ Fe(II) levels in these reactions were monitored for approximately 30 minutes with no ascorbate present, then 2 mM ascorbate was spiked into the reactions and Fe(II) levels measured for another 30 minutes. The resulting data (**Figure 5a**) show a clear decrease in Fe(II) levels in the first 30 minutes of the reaction before ascorbate was added, where reaction mixtures with 2-OG and RNA substrate produced the largest decreases in Fe(II). At 30 minutes, ascorbate (2 mM) was spiked into the reaction mixture and samples thereafter show a marked increase in measured Fe(II) levels and relative stabilization of Fe(II) levels as the reaction proceeds. Additionally, samples from an analogous reaction were monitored for adenosine production by UHPLC-MS before and after an ascorbate spike (**Figure 5b**), showing that ascorbate’s ability to rescue m^6^A demethylation activity and adenosine production is directly correlated to the stabilization of Fe(II) levels during the FTO-catalyzed demethylation reaction.

**Figure 5.**
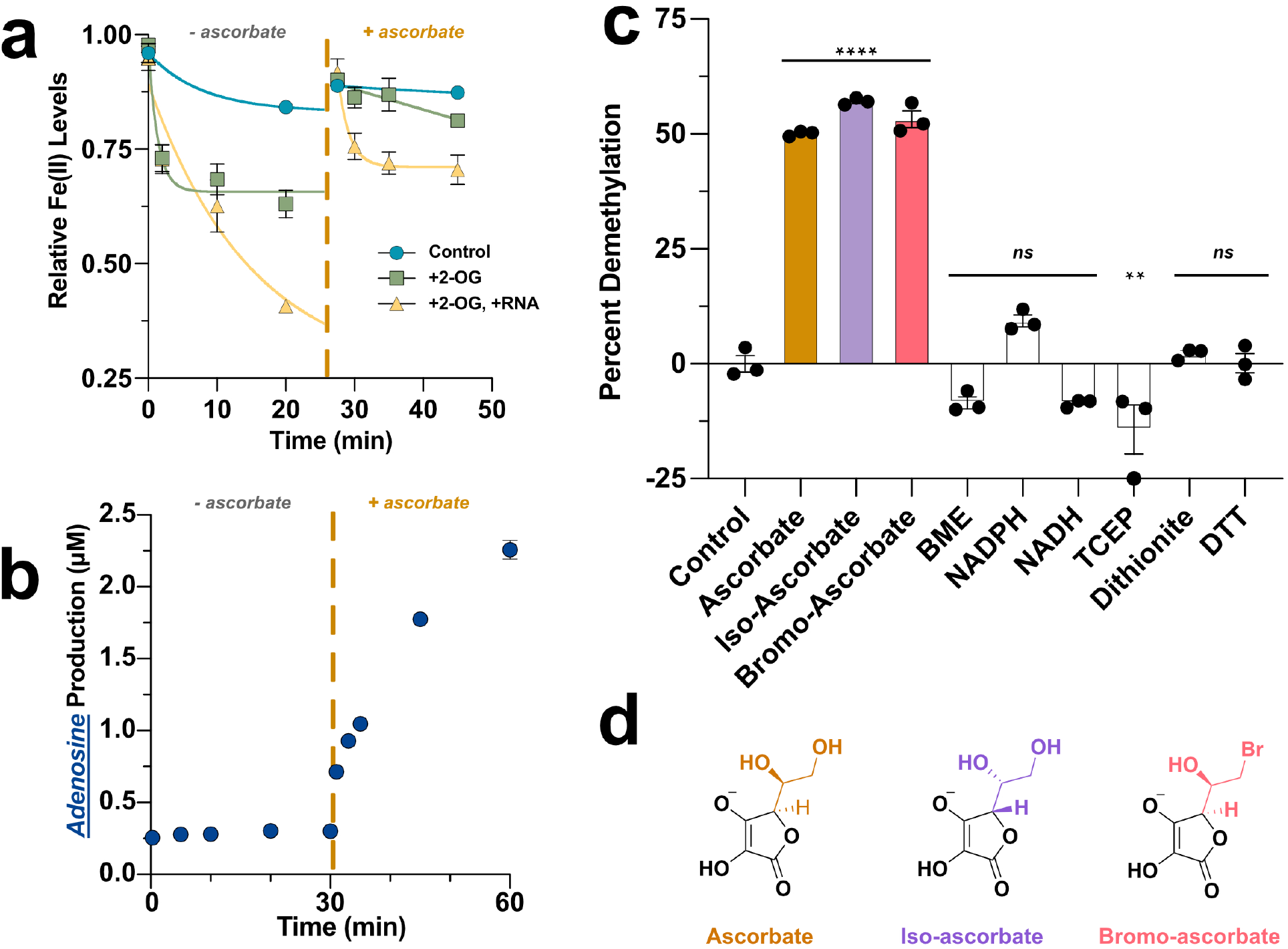
Ascorbate rescues Fe(II) levels and catalytic activity in demethylation reactions with FTO. **(a)** Reactions with FTO, Fe(II), and no ascorbate were set up with either no additional cofactors or substrate (‘control’, blue circles), added 2-OG cofactor (‘+2-OG’, green squares), or added 2-OG and linear 9-mer m6A-RNA substrate (‘+2OG, +RNA’, yellow triangles). Reactions were quenched at different timepoints with phenanthroline and then A_511_ was measured on a plate reader to quantify relative Fe(II) levels. Dotted line indicates the addition of ascorbate into all reaction mixtures to a final concentration of 2 mM. **(b)** Similar +2-OG, +RNA reaction as in **a**, with ascorbate spike-in occuring at the dotted line, where samples were quenched with EDTA and analyzed by UHPLC-MS to quantify adenosine product formation over the course of the reaction. **(c)** End-point reactions of FTO with m^6^A linear 9-mer measuring percent demethylation by UHPLC-MS with 1 mM of various reducing agents. Statistical significance compared to the no reducing agent control was calculated using a one-way ANOVA with Tukey’s multiple comparison test (****p<0.0001, **p<0.005, ns p>0.05). **(d)** Chemical structures of ascorbate and its derivatives iso-ascorbate and bromo-ascorbate.

Prior studies with FODDs CP4H and Tet2 have shown that other reducing agents are unable to act in the same manner as ascorbate to promote activity,^53,56^ suggesting a specific interaction between ascorbate and FODDs. We similarly tested ascorbate selectivity with FTO by running end-point demethylation reactions with m^6^A linear 9-mer substrate against a panel of standard reducing agents, analyzing percent demethylation by UHPLC-MS as above (**Figure 5c**). Non-ascorbate reducing agents showed negligible demethylation activity with FTO, similar to a no reducing agent control, whereas ascorbate, and derivatives iso-ascorbate or bromo-ascorbate (**Figure 5d**), all produced a significant increase in FTO activity (*p<0*.*0001)*. These experiments demonstrate that ascorbate-mediated activation of demethylation by FTO is highly specific, likely because ascorbate must interact directly with the active site of FTO to reduce stalled Fe(III) states.

### Co-crystal structure reveals how ascorbate selectively engages FTO’s active site

Although it has been known for over 50 years that some FODDs are critically dependent on ascorbate, no structures of ascorbate bound to any FODD have been reported and there is scarce biophysical information about how this cofactor selectively engages the FODD active site. Because FTO is highly dependent on ascorbate, we reasoned that this enzyme would be a good model system to capture an FODD-ascorbate interaction. Using Ga(III) to mimic a stalled Fe(III) state, we determined a 3.07 Å co-crystal structure of a truncated FTO construct in complex with ascorbate and Ga(III) (**Figure 6, Supplementary Figure 3, Supplementary Table 2**). The initial structure solution and refinement showed the conserved His-Asp-His triad coordinating the Ga(III) metal center, in addition to significant unmodeled electron density directly adjacent to the metal binding site (**Figure 6a**). Ascorbate fits well within this *F*_o_-*F*_c_ density and refines to a position where both enediol oxygen atoms coordinate the Ga(III) center and would be poised to carry out reduction of Fe(III) formed during uncoupled reaction cycles. The lactone oxygen atom in ascorbate makes a hydrogen bonding interaction with FTO N205 (**Figure 6b**), an active site asparagine residue conserved across many other AlkBH family members (*e*.*g*. AlkB, AlkBH3, AlkBH5). The di-hydroxy ‘arm’ of ascorbate points away from the 2-OG binding pocket and has potential hydrogen bonding contacts with FTO residues R96 and R322. Although the ascorbate arm points away from the 2-OG binding pocket, either 2-OG or succinate binding to the active site metal would overlap with bound ascorbate, suggesting that 2-OG/succinate binding is mutually exclusive with ascorbate binding (**Figure 6c**).

**Figure 6.**
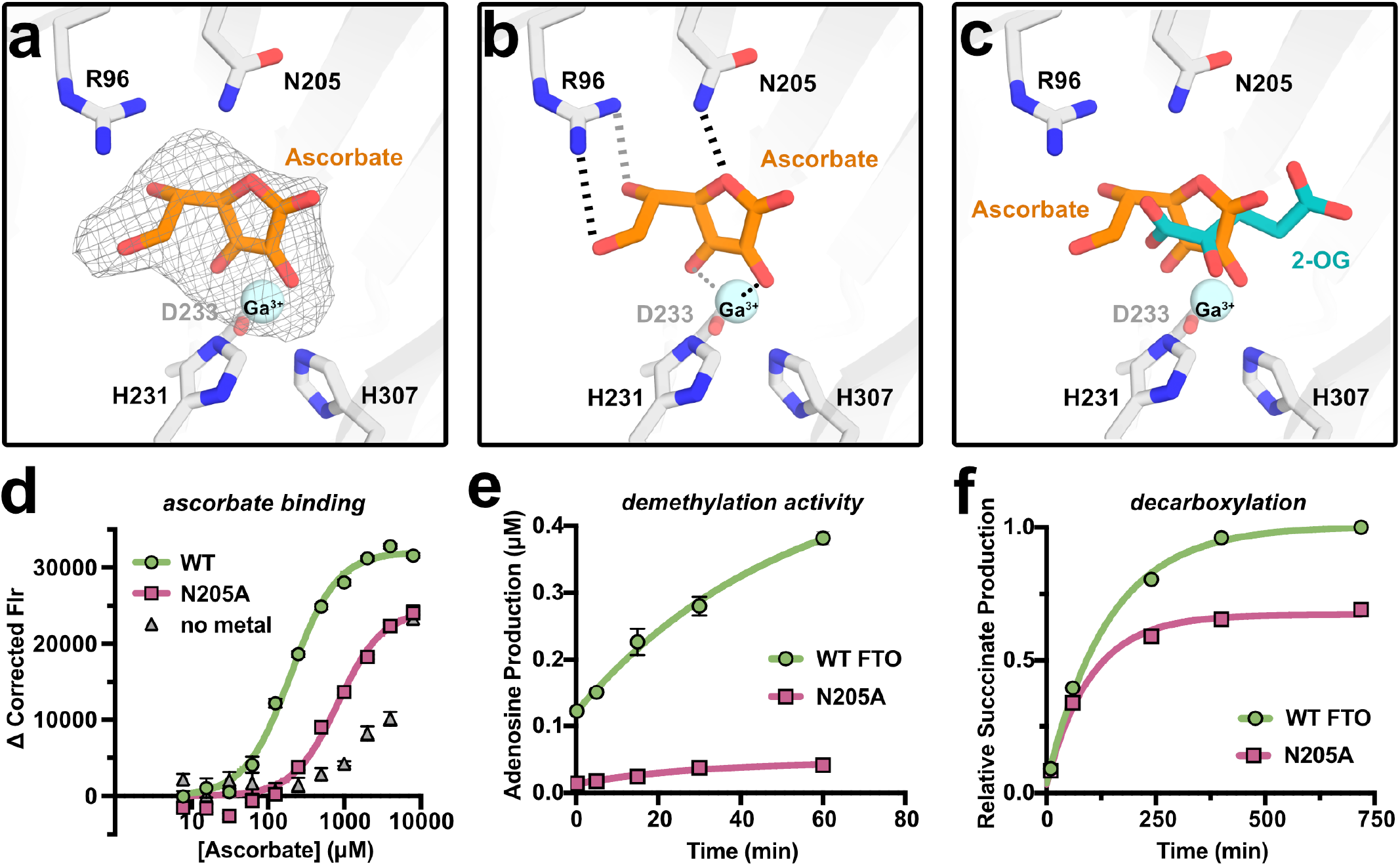
Crystal structure of ascorbate bound to FTO active site reveals details of cofactor recruitment. **(a)** Co-crystal structure of FTO in complex with Ga(III) and ascorbate showing *F*_*O*_*-F*_*C*_ omit map for ascorbate at 3.0σ in the FTO active site. (**b)** Ga(III) is coordinated by the canonical HDH triad and the enediol moiety of ascorbate. Dotted lines show ascorbate interaction with Ga(III) and hashed lines show hydrogen bonding interactions between ascorbate and FTO residues. **(c)** FTO-ascorbate structure with 2-OG aligned from PDB 4ZS2. Ascorbate and 2-OG bind in different, but overlapping positions, suggesting these cofactors (and likely also succinate) cannot bind FTO at the same time. **(d)** Intrinsic tryptophan fluorescence titrations showing ascorbate binding to WT FTO + Ga(III) (green circles), N205A FTO + Ga(III) (pink squares), or WT FTO with no added metal (gray triangles). **Supplementary Table 3** lists the fitted *K*_D_ values for ascorbate binding to WT and N205A FTO; in the absence of metal, ascorbate binding is much weaker and could not be fit to a *K*_D_. **(e)** FTO WT or N205A demethylation reactions with m^6^A-RNA show N205A significantly impairs FTO-mediated demethylation activity. Adenosine product formation over time was measured by UHPLC-MS. **(f)** FTO WT or N205A reactions produce similar amounts of succinate over time in the absence of prime m^6^A-RNA substrate, suggesting N205A minimally affects FTO-mediated 2-OG decarboxylation. Succinate production was measured by ^1^H-NMR in reaction mixures with Fe(II), 2-OG, and ascorbate, but no prime RNA substrate.

### FTO N205A mutation impairs ascorbate binding and demethylation catalysis

To further validate our structural model of the ascorbate-FTO interaction, we used intrinsic tryptophan fluorescence to examine how mutating FTO N205, which hydrogen bonds with ascorbate in our structure, impacts ascorbate binding affinity to FTO. Titrating ascorbate into solutions of Ga(III)-bound FTO WT versus N205A reveals that mutation of N205 reduces ascorbate binding affinity by approximately 4-fold (**Figure 6d, Supplementary Figure 4a**,**b, Supplementary Table 3**). In the absence of active site metal, ascorbate binding was further impaired, and binding curves could not be fit to obtain a *K*_D_. Together, these binding data suggest both FTO N205 and active site metal play important roles in ascorbate recruitment to the FTO active site. Fluorescence quenching experiments were carried out with Ga(III) metal in the FTO active site to mimic the stalled Fe(III) state, but we also observed similar effects with Mn(II) (**Supplementary Figure 4c, Supplementary Table 3**). Next, we compared FTO WT and N205A demethylation activity on an m^6^A-9mer RNA substrate and found that N205A FTO had significantly reduced demethylation activity compared to WT (**Figure 6e**). While the loss of m^6^A-RNA demethylation activity for FTO N205A is consistent with impaired ascorbate binding and the strict ascorbate requirements for this substrate, N205 also makes contacts with 2-OG in the active site (PDB 4ZS2), so this mutation could also impact 2-OG binding and decarboxylation. To distinguish between the possible effects of N205A mutation on ascorbate versus 2-OG interaction, we performed decarboxylation reactions and monitored succinate production via ^1^H-NMR as described above with WT versus N205A FTO. The resulting data show that N205A can still readily decarboxylate 2-OG, at approximately 70 % the level of WT FTO (**Figure 6f)**. These data, along with the fact that N205 is located relatively far from the prime substrate itself (> 6 Å away, PDB 5ZMD), support the conclusion that the loss of m^6^A demethylation activity in the N205A mutant arises at least in part from disruption of ascorbate binding in the FTO active site.

## DISCUSSION

The FODD superfamily of enzymes carries out essential oxidation reactions on a wide range of biological substrates – including peptides, DNA, and RNA – to modify and modulate the structure and function of these biomolecules in the cell. While it has been known for decades that these enzymes frequently rely on cofactor ascorbate for efficient catalysis, we still have a poor understanding of how ascorbate usage varies across this superfamily, how ascorbate usage might change under different conditions or with different substrates, and how ascorbate selectively engages the FODD active site. We compared ascorbate usage for model FODD enzymes FTO and AlkBH5 and surprisingly found that these two closely related RNA demethylases have significantly different ascorbate dependencies for their m^6^A-RNA demethylation reactions (**Figure 2**). Likely related to this, we also show that FTO will readily catalyze the decarboxylation reaction of 2-OG in the absence of prime substrate, whereas AlkBH5 will not (**Figure 4**). Finally, comparing FTO-mediated demethylation of a series of different m^6^A- or m^6^A_m_-containing RNAs, we found that ascorbate usage varies dramatically across these different substrates (**Figure 3**), where prime substrates that are efficiently turned over (higher *k*_cat_) are significantly less dependent on ascorbate than substrates that have slower catalytic steps (lower *k*_cat_).

Together, our biochemical data for RNA demethylases FTO and AlkBH5 suggest two key principles of FODD catalysis that control ascorbate dependence: (1) the propensity of the enzyme to carry out decarboxylation in the absence of substrate (*i*.*e*. uncoupled decarboxylation), and (2) the efficiency of prime substrate catalysis (*i*.*e. k*_cat_). Because FTO readily catalyzes decarboxylation of 2-OG even in the absence of substrate (**Figure 4b,ii**), it likely has a high propensity to enter uncoupled reaction cycles when acting on relatively poor prime substrates (*e*.*g*. m^6^A-RNA with *k*_cat_ = 0.02 min^-1^), giving rise to a strong dependence on ascorbate in order to rescue stalled Fe(III) states. In contrast, because AlkBH5 does not catalyze 2-OG decarboxylation in the absence of prime substrate (**Figure 4b,iii**), it is likely much more resistant to entering uncoupled reaction cycles, making this enzyme much less reliant on ascorbate. However, even though FTO has a high predisposition for uncoupled decarboxylation reactions, with a prime substrate that undergoes rapid or efficient catalysis (*e*.*g*. m^6^A_m_-RNA with *k*_cat_ = 4.7 min^-1^), the prime substrate reaction is apparently able to ‘keep up’ with the decarboxylation reaction and maintain a coupled reaction cycle, dramatically reducing FTO’s ascorbate dependence. Therefore, we propose that the relative rates of uncoupled decarboxylation and prime substrate reactions are key determining features of FODD catalysis that can significantly tune FODD activity and selectivity at different ascorbate concentrations.

Recent biophysical work that combines 2D NMR experiments and MD simulations on both AlkBH5^70^ and FTO^71^ has suggested that the binding of prime, methylated RNA substrate induces a dynamic shift in the active site that rearranges 2-OG binding to the iron center, enabling O_2_ activation and initiating catalysis of both the decarboxylation and hydroxylation reactions. This dynamical model that prevents oxygen activation in the AlkBH5 active site in the absence of prime substrate nicely agrees with our biochemical data showing that AlkBH5 does not readily undergo 2-OG decarboxylation without RNA. However, while a similar dynamic model that requires prime substrate for O_2_ activation and 2-OG decarboxylation was proposed for FTO,^71^ our experimental data show the opposite – that FTO, in contrast to AlkBH5, readily undergoes 2-OG decarboxylation in the absence of prime substrate (**Figure 4**). This suggests that the FTO active site is capable of achieving reactive configurations of Fe-bound 2-OG that allow for efficient oxygen activation and conversion of 2-OG to succinate even the absence of prime substrate, which we argue in large part accounts for FTO’s ascorbate requirement. It is therefore likely that subtle differences in enzyme active site configuration and dynamics across the FODDs will tune how readily different superfamily members undergo oxygen activation and decarboxylation, directly impacting their dependence on ascorbate and overall activity and selectivity.

Finally, we also determined the first structure of ascorbate bound to an FODD active site, using FTO with Ga(III) to mimic a stalled Fe(III) state (**Figure 6**). Our FTO-ascorbate structure suggests that key contacts and complementarity between the ascorbate lactone scaffold and FTO active site – namely metal chelation and hydrogen bonding with N205 – mediate specificity for ascorbate. In contrast, the di-hydroxy arm of ascorbate points away from the active site pocket, consistent with arm substitutions having no effect on ascorbate’s ability to promote FTO-mediated demethylation catalysis (**Figure 5c,d**). Our structural model is further validated by biophysical and biochemical data that show mutating FTO N205 weakens ascorbate binding and selectively impairs catalysis of prime hydroxylation more than decarboxylation (**Figure 6d-f**), suggesting that the N205A mutation does not significantly disrupt 2-OG binding and oxygen activation. The positioning of ascorbate in our structural model overlaps with the 2-OG binding site found in other published structures,^64,72^ suggesting that ascorbate binding is mutually exclusive with 2-OG and succinate binding to the iron center in the active site, and that these molecules must be displaced from the active site before ascorbate can bind and carry out reduction of Fe(III). Furthermore, ascorbate binding in FTO differs from CMD1, an algal dioxygenase that uses ascorbate as a co-substrate in lieu of 2-OG to modify DNA.^73^ In CMD1, ascorbate coordinates the metal center in a monodentate manner,^73^ and the arm of ascorbate extends into the CMD1 active site pocket. In contrast, our model supports bidentate metal chelation, similar to the way other enediol moieties interact with enzyme metal centers.^74,75^ Together our structure and accompanying biochemical data provide the first atomic-level model of how ascorbate binds FODD active sites and identifies key interactions that mediate ascorbate binding specificity.

Ascorbate concentrations vary widely across different human cell types and disease states – from 0.2 to 20 mM.^52,76^ Our mechanistic framework for the differential, ascorbate-dependent control of FTO and AlkBH5 activities therefore suggests that cellular m^6^A RNA demethylation across the transcriptome will be selectively tuned by the ascorbate concentration in different environments. Consistent with this, a recent study showed that global m^6^A levels decrease in HEK293T cells with the addition of excess exogenous ascorbate.^51^ In analogy to FTO and AlkBH5, we propose that other AlkBH family demethylases, and the FODD superfamily more broadly, will have a wide range of ascorbate-dependent activity that is controlled by the relative kinetics of an individual enzyme’s intrinsic decarboxylation reaction and the efficiency of prime substrate catalysis. In this way, cellular ascorbate concentration likely plays a thus far underappreciated but important role in selectively regulating RNA, DNA, and polypeptide oxidation reactions across the cell and in diverse human diseases.

## METHODS

### FTO and AlkBH5 vector construction

The human FTO(32-505) DNA sequence was obtained from IDT as a codon-optimized gblock and cloned into a pET28a bacterial expression vector with N-terminal 6xHis tag. The FTO N205A single point mutation was introduced into the 6xHis-FTO(32-505) construct using whole plasmid PCR site-directed mutagenesis. FTO(Δ166-189) is an alteration of 6xHis-FTO(32-505) that deletes flexible loop residues 166-189, prepared using whole plasmid PCR with non-adjacent primers. The human AlkBH5(66-292) DNA sequence was obtained from IDT as a codon-optimized gblock and cloned into a pET28a bacterial expression vector with N-terminal 6xHis tag and tobacco etch virus (TEV) protease cleavage site (ENLYFQGS) located between the 6xHis tag and AlkBH5 protein.

### FTO and AlkBH5 overexpression

All proteins were all overexpressed in *E. coli* cells as follows: bacterial expression constructs were transformed into *E. coli* BL21 (DE3) Rosetta cells, plated on LB agar plates with Kanamycin (50 μg/mL) and Chloramphenicol (25 μg/mL). Seed cultures were grown overnight from single colonies in LB media at 37 °C with shaking at 200 rpm. Seed cultures were used to inoculate primary cultures (1:100 dilution, 37 °C, 200 rpm) and once OD_600_ of 0.8 was reached, protein expression was induced with 1 mM isopropyl β-D-1-thiogalactopyranoside (IPTG) at 16 °C overnight (∼18 hours, 200 rpm). Cells were pelleted via centrifugation (7500 x *g* for 45 minutes), the supernatant was discarded, and the pellets were flash frozen and stored at -70 °C.

### FTO purification

FTO(32-505), FTO(N205A), and FTO(Δ-166-189) were all purified following the same procedure. Frozen FTO cell pellets were thawed and resuspended in lysis buffer (25 mM Tris pH 7.5, 300 mM NaCl, 10 mM imidazole, 0.05% NP40, 1 mM BME), sonicated, then lysates were clarified via centrifugation (23000 x *g* for 45 minutes). Supernatant was loaded onto NiNTA resin (5 mL bed volume, 1 hour batch binding) in a gravity column, washed with 25 mL wash buffer (25 mM Tris pH 7.5, 500 mM NaCl, 40 mM imidazole, 1 mM BME) and eluted with 25 mL elution buffer (25 mM Tris pH 7.5, 200 mM NaCl, 400 mM imidazole, 1 mM BME). Eluate was then buffer exchanged using centrifugal filters (30 kDa MWCO) into anion exchange buffer A (25 mM Tris pH 7.5, 10 mM NaCl) before loading onto a GE/Cytiva AKTA Pure FPLC system with Cytiva HiTrap 5mL Q HP column. FTO was eluted from the anion exchange column with a gradient into 100% anion exchange buffer B (25 mM Tris pH 7.5, 750 mM NaCl) over 45 minutes. Fractions with FTO were collected and concentrated before loading onto a Cytiva HiLoad 16/600 Superdex 200 pg gel filtration column, and eluted with an isocratic gradient of 25 mM HEPES pH 7, 50 mM KCl. Pure FTO fractions were collected and concentrated to 200 μM in gel filtration buffer, flash frozen in liquid nitrogen, and stored at -70 °C.

### AlkBH5 purification

AlkBH5(66-292) was purified in a similar manner as FTO, without the ion exchange purification step. Frozen AlkBH5 pellets were lysed in lysis buffer (25 mM HEPES pH 8, 300 mM NaCl, 0.1% Triton X, 10 mM imidazole) via sonication, lysates were clarified via centrifugation (23000 x *g* for 45 minutes), and supernatant was loaded onto a NiNTA gravity column. The resin (5 mL bed volume, 1 hour batch binding) was washed (25 mM HEPES pH 8, 750 mM NaCl, 25 mM imidazole, 25 mL) and eluted with 25 mL elution buffer (25 mM HEPES pH 8, 250 mM NaCl, 400 mM imidazole). The N-terminal 6xHis tag was then removed by incubating overnight with TEV protease (1:50 molar ratio of TEV:protein, 4 °C, 18 hrs). The TEV-cleaved protein was buffer-exchanged with a centrifugal spin filter (10 kDa MWCO) back into lysis buffer for a back-pass through 2 mL bed volume NiNTA resin to remove the cleaved 6xHis tag and TEV protease. The NiNTA back-pass flowthrough was concentrated and loaded onto a Cytiva HiLoad 16/600 Superdex 200 pg column for gel filtration into storage buffer (25 mM HEPES pH 7, 100 mM KCl). Pure AlkBH5 fractions were collected and concentrated to 200 μM in storage buffer, flash frozen in liquid nitrogen, and stored at -70 °C.

### RNA substrates for demethylation assays

m^6^A linear 9-mer RNA (CUGG(m^6^A)CUGG) and m^6^A 25-mer stem-loop RNA (GUUGGGCCUC(m^6^A)GGUUGGCCCUUGG) were purchased from IDT. m^6^A_m_ 5′ cap RNA (m^7^Gppp(m^6^A_m_)CUGG) was synthesized analogously to previously reported m^6^A_m_ cap oligonucleotides.^77^ All substrates were resuspended in water at 500 μM and stored at -70 °C.

### Demethylation assays For Michaelis-Menten kinetics

Reactions were performed with FTO (200 or 500 nM) or AlkBH5 (100 or 200 nM), 2-oxoglutarate (300 μM), (NH_4_)_2_Fe(SO_4_)_2_•6H_2_O (75 μM), L-ascorbic acid (0 or 2 mM), and m^6^A-containing substrate (varying concentrations) in reaction buffer (25 mM HEPES pH 7, 50 mM NaCl, and 50 µM caffeine as MS internal standard). Reactions were performed at 37 °C (FTO) or 30 °C (AlkBH5) for 5 minutes and were initiated by the addition of a 2X substrate solution into a 2X protein and cofactor solution. Note: a lower reaction temperature was required for AlkBH5 here in order to be able to capture initial rates by manual pipetting/quenching; this makes cross-comparison of FTO versus AlkBH5 kinetic parameters difficult, but we rely only on comparison of kinetic parameters across different substrates for FTO, or different substrates for AlKBH5, in this manuscript. Next, reaction timepoints (10 μL) were removed and quenched at 0.25, 1, 2, 3, and 5 minutes by addition of EDTA (1 mM final concentration in quenched timepoint). Quenched reaction samples were digested to single nucleosides with Nucleoside Digestion Mix (NEB, M0694S) per commercial protocol, but with approximately half of the recommended nuclease concentration. Samples with m^6^A_m_ cap substrates were first decapped with mRNA Decapping Enzyme (NEB, M0608S) per commercial protocol for 1 hour, and then digested to single nucleosides as above.

### UHPLC-MS nucleoside analysis

RNase-digested reaction samples were loaded onto an Agilent Bio-Inert 1260 Infinity II UHPLC system with Infinity Lab LC/MSD iQ, equipped with an Agilent Zorbax SB-Aq Rapid Resolution HD (2.1 × 100 mm, 1.8 µm particle size) using mobile phase containing 0.1% formic acid (A) and 100% acetonitrile, 0.1% formic acid (B). Nucleosides were detected and quantified by mass spectrometry in positive ionization mode. The LC gradient at flow rate 0.480 mL/min was as follows for detection of m^6^A (m/z = 282, 3.5 min) and A (m/z = 268, 1.6 min): 0-1.0 min, 100% A; 1.0-4.25 min, to 85% A/15% B; 4.25-5.0min, to 25% A/75% B; 5.0-7.0 min, 25% A/75% B; 7.0-7.1 min, to 100% B; 7.1-11.75 min, 100% B. The LC gradient at flow rate 0.450 mL/min was as follows for detection of m^6^A_m_ (m/z = 296, 3.65 min) and A_m_ (m/z = 282, 3.5 min): 0-1.0 min, 100% A; 1.0-1.1 min, to 40% A/60% B; 1.1-5.0 min, to 28% A/72% B; 5.0-5.1 min, to 100% B; 5.1.0-8.0 min, 100% B. Digested nucleoside peak identity and quantification was obtained by comparing to nucleoside standards (**Supplementary Figure 1a**).

### Michaelis-Menten kinetic analysis

Concentrations of adenosine (or 2′-*O*-methyladenosine) product in quenched and nuclease-digested reaction samples were obtained by comparison of MS peak areas to a linear dilution series of synthetic nucleoside standard (A or A_m_) at known concentrations by UHPLC-MS. Initial rates were calculated by plotting the product A or A_m_ concentration over time and fitting the resulting plot to a linear equation to obtain an observed rate constant (**Supplementary Figure 1b**). Observed rate constants were then plotted against RNA substrate concentration and fit to the standard Michaelis-Menten equation in GraphPad Prism to obtain kinetic parameters.

### FTO ascorbate sensitivity assays with various RNA substrates

Reactions were set up with FTO (1 μM), 2-oxoglutarate (300 μM), (NH_4_)_2_Fe(SO_4_)_2_•6H_2_O (75 μM), 2.5 μM substrate (either m^6^A linear 9-mer, m^6^A stem-loop 25-mer, or m^6^A_m_ cap RNA), and varying levels of ascorbate (0 – 2 mM) in reaction buffer (above). Reactions were initiated by mixing 2X protein and cofactor solution with 2X RNA and run at 30 °C for 1 hour. Time zero and 1 hour endpoint samples were taken and quenched with EDTA (1 mM final concentration). Quenched reaction samples were then digested for 1 hour or overnight with Nucleoside Digestion Mix (NEB) for m^6^A- or m^6^A_m_-containing substrates, respectively. m^6^A and m^6^A_m_ nucleoside levels were quantified by UHPLC-MS as described above. For each substrate, percent demethylation was calculated by dividing the amount of m^6^A or m^6^A_m_ consumed after 1 hour by the amount of m^6^A or m^6^A_m_ at time zero in the 0 mM ascorbate reaction.

### Reducing agent specificity panel

Reaction mixtures were prepared with FTO (2 μM), 2-oxoglutarate (300 μM), (NH_4_)_2_Fe(SO_4_)_2_•6H_2_O (75 μM), 4 μM RNA substrate (m^6^A linear 9-mer), and variable reducing agent (1 mM, see **Figure 5**) or a no reducing agent negative control. Reactions were initiated by mixing 2X protein and cofactor solution with 2X RNA, with endpoint samples taken at 150 minutes and quenched with EDTA (1 mM final). Samples were digested and analyzed via UHPLC-MS as above. Percent demethylation was calculated by comparing the amount of m^6^A measured after 2.5 hours to the amount of m^6^A at time zero in the negative control (no reducing agent) reaction.

### Quantification of Fe(II) levels during FTO reactions

Reaction mixtures were prepared with FTO (2 μM), 2-oxoglutarate (300 μM), and (NH_4_)_2_Fe(SO_4_)_2_ 6H_2_O (15 μM), and linear 9-mer RNA substrate (2.5 μM) in the absence of ascorbate. Reactions were initiated by mixing 2X protein and cofactor solution with 2X RNA. Reactions progressed for 27 minutes and then ascorbate was spiked in to a final concentration of 1 mM ascorbate. For both the pre- and post-ascorbate periods of the reaction, timepoint samples (45 μL) were removed and quenched with phenanthroline (250 μM final concentration). Phenanthroline tightly chelates iron and selectively absorbs at ∼ 511 nm when complexed with Fe(II).^69^ Absorbance at 511 nm of the phenanthroline-quenched timepoint samples was measured on a Tecan Spark Multimode Plate Reader by averaging the absorbance of wavelengths 509-513 nm (which we call ‘A_511_’). A_511_ values were normalized to obtain the relative Fe(II) levels in each phenanthroline-quenched sample over the course of the reaction. An analogous reaction, but quenched with EDTA, was carried out in order to monitor adenosine concentrations by UHPLC-MS. This reaction was setup identically to the one described above and timepoint samples (10 μL) were quenched with EDTA (1 mM final), with an ascorbate spike (1 mM final) at 30-minutes, followed by quantification of adenosine production by UHPLC-MS as above.

### FTO-ascorbate structure determination and refinement

Purified 6xHis-FTO(Δ166-189) (20 mg/mL) was incubated for 30 minutes at room temperature with Ga(NO_3_)_3_ (1 mM) and sodium L-ascorbate (10 mM, prepared fresh) in 10 mM HEPES pH 8, 50 mM NaCl. The protein and cofactor solution was mixed 1:1 with precipitant solution (100 mM MES pH 6.5, 1.6 M ammonium sulfate, 10% (v/v) 1,4-dioxane) using a Formulatrix NT8 dropsetter into SwissSci 3-well Midi 96-well sitting drop trays, with 200 μL drop volumes. Initial crystals were harvested, crushed using Hampton Research Bead-Beater, and seeded into new drops with the NT8 drop setting robot (20 μL seed stock, 180 μL protein mixture, 200 μL precipitant solution). Rhombohedral crystals grew after 1 day, were harvested and flash-frozen in precipitant well solution with 25% glycerol as cryoprotectant, and sent to Cornell High Energy Synchrotron Source (CHESS) for X-ray diffraction experiments (FlexX beamline ID7B2 with Dectris Eiger2 16M detector and collection at 100K). Diffraction data were indexed and processed with XDS. Molecular replacement with PHASER/PHENIX and PDB 6OHS was used to solve the structure of FTO and further refinements and model building were carried out in PHENIX and COOT. After placement of Ga(III) metal and refinement of protein residues, ascorbate was modeled into the large, positive *F*_*O*_*-F*_*C*_ density remaining in the active site using COOT; ascorbate restraints were generated using Grade2 on the Grade Web Server (https://grade.globalphasing.org).^78^ Subsequent rounds of automated refinement and manual adjustments were done with PHENIX and COOT to arrive at the final structural model. Data collection and refinement parameters can be found in **Supplementary Table 2**.

### Intrinsic tryptophan fluorescence assays to quantify ascorbate binding

15 μL solutions of FTO (20 μM final concentration) and metal (Ga(NO_3_)_3_ or MnCl_2_, 40 μM final concentration) with varying concentrations of ascorbate (8000 – 0 μM) in buffer (25 mM HEPES pH 7.5, 50 mM KCl) were added to Grenier 384-well flat-bottom, black, low-volume microplates. The intrinsic tryptophan fluorescence was measured on a Tecan Spark Multimode Plate Reader by exciting FTO’s tryptophan residues at Ex_285nm_ (Gain 140, 10 nm bandwidth) and reading the resulting emission spectra from 316 to 346 nm every 2 nm (5 nm bandwidth). Catalase (at 200 μM concentration to approximately match FTO Trp fluorescence intensity) was used as a control protein to correct for both the inner filter effect and non-specific quenching from high ascorbate concentrations (**Supplementary Figure 4b)**. Using catalase control titrations with ascorbate, a *correction factor* was calculated at each ascorbate concentration (0 - 8 mM):

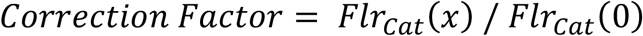

Where *Flr*_*Cat*_*(x)* is the averaged fluorescence intensity of the catalase Trp fluorescence peak maximum (320-328 nm) at a given ascorbate concentration (x), and *Flr*_*Cat*_*(0)* is the averaged fluorescence intensity of the catalase peak maximum at 0 mM ascorbate.

We then used this *correction factor* from the catalase control titration to correct for the inner filter effect and nonspecific binding from high ascorbate concentrations, by applying the following formula to Trp fluorescence data collected in our FTO-ascorbate titration experiments:

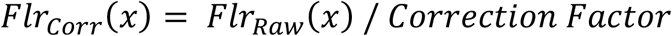

Where *Flr*_*Raw*_*(x)* is the average measured fluorescence intensity of the FTO Trp fluorescence peak maximum (330-338 nm) at a given ascorbate concentration (x), and *Flr*_*Corr*_*(x)* is the resulting corrected fluorescence intensity.

We next calculated the change in corrected FTO fluorescence intensity (Δ Corrected Flr) during the ascorbate titration by subtracting the corrected fluorescence intensity at a given ascorbate concentration (*Flr*_*Corr*_*(x)*) from the corrected fluorescence intensity at 0 mM ascorbate (*Flr*_*Corr*_*(0)*):

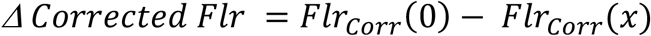

The difference in corrected fluorescence values over the course of the titration (‘Δ Corrected Flr’ in **Figure 6d**) were plotted versus ascorbate concentration and fit using a Hill binding model in GraphPad Prism to obtain *K*_*D*_ values.

### ^1^H NMR assays to monitor decarboxylation reactions

A reaction mixture with 2-OG (100 μM), ascorbate (500 μM), (NH_4_)_2_Fe(SO_4_)_2_•6H_2_O (25 μM), and 3-(Trimethylsilyl)propionic-2,2,3,3-d4 (TSP; NMR internal standard) was prepared in 0.5X PBS with 90% H_2_O / 10% D_2_O, and either FTO or AlkBH5 (10 μM). Reactions were initiated with the addition of enzyme and ^1^H NMR spectra were measured (using excitation sculpting scheme for water suppression) approximately every hour after an initial acquisition with a 600 MHz Bruker NMR spectrometer equipped a 5-mm QCI cryoprobe. All spectra were collected at 298 K with 64 scans, 4 dummy scans, and a delay time of 5 seconds. Data was processed using Topspin using automatic phase- and baseline-correction. The singlet at 2.32 ppm (succinate) and triplet at 2.36 ppm (2-OG) were integrated manually and normalized to the TSP internal standard to quantitatively monitor relative changes in succinate and 2-OG levels over time.

## ACKNOWLEDGMENTS

We would like to thank Dr. Shi Bai for help with NMR experiments.

## AUTHOR CONTRIBUTIONS

Conceptualization and Supervision: JSM. Methodology: LOC, JSM. Resources: LOC, MW, JK, JJ. Investigation: LOC. Visualization: LOC, JSM. Writing and editing: LOC, JSM. Funding Acquisition: JSM

## DATA AVAILABILITY

Coordinates and structure factors for the FTO-ascorbate crystal structure were deposited in the Protein Data Bank with accession code 9OHS.

## SUPPLEMENTARY MATERIALS

Supplementary Figures and Tables can be found in the separate Supplementary Materials file.

## REFERENCES

(1) Schofield, C. J.. Z. Z. 2-Oxoglutarate-Dependent Oxygenases and Related Enzymes. Curr. Opin. Struct. Biol. 1999, 9 (6), 722–731.

(2) He, C.; Mishina, Y. Oxidative Dealkylation DNA Repair Mediated by the Mononuclear Non-Heme Iron AlkB Proteins. J. Inorg. Biochem. 2006, 100 (4), 670–678.

(3) Islam, S.; Leissing, T. M.; Chowdhury, R.; Hopkinson, R. J.; Schofield, C. J. 2-Oxoglutarate-Dependent Oxygenases. 2018, 585–620.

(4) Herr, C. Q.; Hausinger, R. P. Amazing Diversity in Biochemical Roles of Fe(II)/2-Oxoglutarate Oxygenases. Trends Biochem. Sci. 2018, 43 (7), 517–532. 10.1016/j.tibs.2018.04.002.

(5) Pinnell, S. R. Regulation of Collagen Biosynthesis by Ascorbic Acid: A Review. Yale J. Biol. Med. 1985, 58 (6), 553–559.

(6) Peterkofsky, B. Ascorbate Requirement for Hydroxylation and Secretion of Procollagen: Relationship to Inhibition of Collagen Synthesis in Scurvy. Am. J. Clin. Nutr. 1991, 54 (SUPPL. 6), 1135S–1140S. 10.1093/ajcn/54.6.1135s.

(7) Nelson, P. J.; Pruitt, R. E.; Henderson, L. L.; Jenness, R.; Henderson, L. M. Effect of Ascorbic Acid Deficiency on the In Vivo Synthesis of Carnitine. 1981, 672, 123–127.

(8) Vaz, F. M.; van Vlies, N. Dioxygenases of Carnitine Biosynthesis: 6-N-Trimethyllysine and γ-Butyrobetaine Hydroxylases. In 2-Oxoglutarate-Dependent Oxygenases; Schofield, C., Hausinger, R., Eds.; The Royal Society of Chemistry, 2015; p 0. 10.1039/9781782621959-00324.

(9) Schofield, C. J.; Ratcliffe, P. J. Oxygen Sensing by HIF Hydroxylases. Nat. Rev. Mol. Cell Biol. 2004, 5 (5), 343–354. 10.1038/nrm1366.

(10) Elvidge, G. P.; Glenny, L.; Appelhoff, R. J.; Ratcliffe, P. J.; Ragoussis, J.; Gleadle, J. M. Concordant Regulation of Gene Expression by Hypoxia and 2-Oxoglutarate-Dependent Dioxygenase Inhibition: The Role of HIF-1α, HIF-2α, and Other Pathways. J. Biol. Chem. 2006, 281 (22), 15215–15226. 10.1074/jbc.M511408200.

(11) Kuiper, C.; Dachs, G. U.; Munn, D.; Currie, M. J.; Robinson, B. A.; Pearson, J. F.; Vissers, M. C. M. Increased Tumor Ascorbate Is Associated with Extended Disease-Free Survival and Decreased Hypoxia-Inducible Factor-1 Activation in Human Colorectal Cancer. Front. Oncol. 2014, 4 FEB (February), 1–10. 10.3389/fonc.2014.00010.

(12) Duncan, T.; Trewick, S. C.; Koivisto, P.; Bates, P. A.; Lindahl, T.; Sedgwick, B. Reversal of DNA Alkylation Damage by Two Human Dioxygenases. Proc. Natl. Acad. Sci. U. S. A. 2002, 99 (26), 16660–16665. 10.1073/pnas.262589799.

(13) Aas, P. A.; Otterlei, M.; Falnes, P.; Vågbø, C. B.; Skorpen, F.; Akbari, M.; Sundheim, O.; Bjørås, M.; Slupphaug, G.; Seeberg, E.; Krokan, H. E. Human and Bacterial Oxidative Demethylases Repair Alkylation Damage in Both RNA and DNA. Nature 2003, 421 (6925), 859–863. 10.1038/nature01363.

(14) Holland, P. J.; Hollis, T. Structural and Mutational Analysis of Escherichia Coli AlkB Provides Insight into Substrate Specificity and DNA Damage Searching. PLoS One 2010, 5 (1). 10.1371/journal.pone.0008680.

(15) Tahiliani, M.; Koh, K. P.; Shen, Y.; Pastor, W. A.; Bandukwala, H.; Brudno, Y.; Agarwal, S.; Iyer, L. M.; Liu, D. R.; Aravind, L.; Rao, A. Conversion of 5-Methylcytosine to 5-Hydroxymethylcytosine in Mammalian DNA by MLL Partner TET1. Science (80-.). 2009, 324 (5929), 930–935. 10.1126/science.1170116.

(16) An, J.; Rao, A.; Ko, M. TET Family Dioxygenases and DNA Demethylation in Stem Cells and Cancers. Exp. Mol. Med. 2017, 49 (4). 10.1038/emm.2017.5.

(17) Feng, Y.; Li, X.; Cassady, K.; Zou, Z.; Zhang, X. TET2 Function in Hematopoietic Malignancies, Immune Regulation, and DNA Repair. Front. Oncol. 2019, 9 (APR), 1–9. 10.3389/fonc.2019.00210.

(18) Bian, K.; Lenz, S. A. P.; Tang, Q.; Chen, F.; Qi, R.; Jost, M.; Drennan, C. L.; Essigmann, J. M.; Wetmore, S. D.; Li, D. DNA Repair Enzymes ALKBH2, ALKBH3, and AlkB Oxidize 5-Methylcytosine to 5-Hydroxymethylcytosine, 5-Formylcytosine and 5-Carboxylcytosine in Vitro. Nucleic Acids Res. 2019, 47 (11), 5522–5529. 10.1093/nar/gkz395.

(19) Kuznetsov, N. A.; Kanazhevskaya, L. Y.; Fedorova, O. S. DNA Demethylation in the Processes of Repair and Epigenetic Regulation Performed by 2-Ketoglutarate-Dependent DNA Dioxygenases. Int. J. Mol. Sci. 2021, 22 (19). 10.3390/ijms221910540.

(20) Roundtree, I. A.; Evans, M. E.; Pan, T.; He, C. Dynamic RNA Modifications in Gene Expression Regulation. Cell 2017, 169 (7), 1187–1200. 10.1016/j.cell.2017.05.045.

(21) Zaccara, S.; Ries, R. J.; Jaffrey, S. R. Reading, Writing and Erasing MRNA Methylation. Nat. Rev. Mol. Cell Biol. 2019, 20 (10), 608–624. 10.1038/s41580-019-0168-5.

(22) Flamand, M. N.; Tegowski, M.; Meyer, K. D. The Proteins of MRNA Modification: Writers, Readers, and Erasers. Annu. Rev. Biochem. 2023, 92, 145–173. 10.1146/annurev-biochem-052521-035330.

(23) Kuiper, C.; Vissers, M. C. M. Ascorbate as a Cofactor for Fe-and 2-Oxoglutarate Dependent Dioxygenases: Physiological Activity in Tumour Growth and Progression. Front. Oncol. 2014, 4 (NOV), 1–11. 10.3389/fonc.2014.00359.

(24) Fitzsimmons, C. M.; Batista, P. J. It’s Complicated… m 6 A-Dependent Regulation of Gene Expression in Cancer. Biochim. Biophys. Acta - Gene Regul. Mech. 2019, 1862 (3), 382–393. 10.1016/j.bbagrm.2018.09.010.

(25) Thomas, J. M.; Batista, P. J.; Meier, J. L. Metabolic Regulation of the Epitranscriptome. ACS Chem. Biol. 2019, 14 (3), 316–324. 10.1021/acschembio.8b00951.

(26) Losman, J. A.; Koivunen, P.; Kaelin, W. G. 2-Oxoglutarate-Dependent Dioxygenases in Cancer. Nat. Rev. Cancer 2020, 20 (12), 710–726. 10.1038/s41568-020-00303-3.

(27) Wong, S. H. K.; Goode, D. L.; Iwasaki, M.; Wei, M. C.; Kuo, H. P.; Zhu, L.; Schneidawind, D.; Duque-Afonso, J.; Weng, Z.; Cleary, M. L. The H3K4-Methyl Epigenome Regulates Leukemia Stem Cell Oncogenic Potential. Cancer Cell 2015, 28 (2), 198–209. 10.1016/j.ccell.2015.06.003.

(28) Li, Z.; Weng, H.; Su, R.; Weng, X.; Zuo, Z.; Li, C.; Huang, H.; Nachtergaele, S.; Dong, L.; Hu, C.; Qin, X.; Tang, L.; Wang, Y.; Hong, G. M.; Huang, H.; Wang, X.; Chen, P.; Gurbuxani, S.; Arnovitz, S.; Li, Y.; Li, S.; Strong, J.; Neilly, M. B.; Larson, R. A.; Jiang, X.; Zhang, P.; Jin, J.; He, C.; Chen, J. FTO Plays an Oncogenic Role in Acute Myeloid Leukemia as a N6-Methyladenosine RNA Demethylase. Cancer Cell 2017, 31 (1), 127–141. 10.1016/j.ccell.2016.11.017.

(29) Cui, Q.; Shi, H.; Ye, P.; Li, L.; Qu, Q.; Sun, G.; Sun, G.; Lu, Z.; Huang, Y.; Yang, C. G.; Riggs, A. D.; He, C.; Shi, Y. M6A RNA Methylation Regulates the Self-Renewal and Tumorigenesis of Glioblastoma Stem Cells. Cell Rep. 2017, 18 (11), 2622–2634. 10.1016/j.celrep.2017.02.059.

(30) Crake, R. L. I.; Burgess, E. R.; Royds, J. A.; Phillips, E.; Vissers, M. C. M.; Dachs, G. U. The Role of 2-Oxoglutarate Dependent Dioxygenases in Gliomas and Glioblastomas: A Review of Epigenetic Reprogramming and Hypoxic Response. Front. Oncol. 2021, 11 (March), 1–17. 10.3389/fonc.2021.619300.

(31) Brągiel-Pieczonka, A.; Lipka, G.; Stapińska-Syniec, A.; Czyżewski, M.; Żybura-Broda, K.; Sobstyl, M.; Rylski, M.; Grabiec, M. The Profiles of Tet-Mediated DNA Hydroxymethylation in Human Gliomas. Front. Oncol. 2022, 12 (April), 1–9. 10.3389/fonc.2022.621460.

(32) Lei, X.; Xu, J. F.; Chang, R. M.; Fang, F.; Zuo, C. H.; Yang, L. Y. JARID2 Promotes Invasion and Metastasis of Hepatocellular Carcinoma by Facilitating Epithelial-Mesenchymal Transition through PTEN/AKT Signaling. Oncotarget 2016, 7 (26), 40266–40284. 10.18632/oncotarget.9733.

(33) Wagner, K. W.; Alam, H.; Dhar, S. S.; Giri, U.; Li, N.; Wei, Y.; Giri, D.; Cascone, T.; Kim, J. H.; Ye, Y.; Multani, A. S.; Chan, C. H.; Erez, B.; Saigal, B.; Chung, J.; Lin, H. K.; Wu, X.; Hung, M. C.; Heymach, J. V.; Lee, M. G. Kdm2a Promotes Lung Tumorigenesis by Epigenetically Enhancing Erk1/2 Signaling. J. Clin. Invest. 2013, 123 (12), 5231–5246. 10.1172/JCI68642.

(34) Shi, L.; Sun, L.; Li, Q.; Liang, J.; Yu, W.; Yi, X.; Yang, X.; Li, Y.; Han, X.; Zhang, Y.; Xuan, C.; Yao, Z.; Shang, Y. Histone Demethylase JMJD2B Coordinates H3K4/H3K9 Methylation and Promotes Hormonally Responsive Breast Carcinogenesis. Proc. Natl. Acad. Sci. U. S. A. 2011, 108 (18), 7541–7546. 10.1073/pnas.1017374108.

(35) Kim, J. H.; Sharma, A.; Dhar, S. S.; Lee, S. H.; Gu, B.; Chan, C. H.; Lin, H. K.; Lee, M. G. UTX and MLL4 Coordinately Regulate Transcriptional Programs for Cell Proliferation and Invasiveness in Breast Cancer Cells. Cancer Res. 2014, 74 (6), 1705–1717. 10.1158/0008-5472.CAN-13-1896.

(36) Aprelikova, O.; Chen, K.; El Touny, L. H.; Brignatz-Guittard, C.; Han, J.; Qiu, T.; Yang, H. H.; Lee, M. P.; Zhu, M.; Green, J. E. The Epigenetic Modifier JMJD6 Is Amplified in Mammary Tumors and Cooperates with C-Myc to Enhance Cellular Transformation, Tumor Progression, and Metastasis. Clin. Epigenetics 2016, 8 (1), 1–16. 10.1186/s13148-016-0205-6.

(37) Li, L.; Zang, L.; Zhang, F.; Chen, J.; Shen, H.; Shu, L.; Liang, F.; Feng, C.; Chen, D.; Tao, H.; Xu, T.; Li, Z.; Kang, Y.; Wu, H.; Tang, L.; Zhang, P.; Jin, P.; Shu, Q.; Li, X. Fat Mass and Obesity-Associated (FTO) Protein Regulates Adult Neurogenesis. Hum. Mol. Genet. 2017, 26 (13), 2398–2411. 10.1093/hmg/ddx128.

(38) Liu, H.; Xie, Y.; Wang, X.; Abboud, M. I.; Ma, C.; Ge, W.; Schofield, C. J. Exploring Links between 2-Oxoglutarate-Dependent Oxygenases and Alzheimer’s Disease. Alzheimer’s Dement. 2022, 18 (12), 2637–2668. 10.1002/alz.12733.

(39) Johnston, C. S.; Corte, C.; Swan, P. D. Marginal Vitamin C Status Is Associated with Reduced Fat Oxidation during Submaximal Exercise in Young Adults. Nutr. Metab. 2006, 3, 1–5. 10.1186/1743-7075-3-35.

(40) Green, H. L. H.; Brewer, A. C. Dysregulation of 2-Oxoglutarate-Dependent Dioxygenases by Hyperglycaemia: Does This Link Diabetes and Vascular Disease? Clin. Epigenetics 2020, 12 (1), 1–15. 10.1186/s13148-020-00848-y.

(41) Yang, Z.; Yu, G. li; Zhu, X.; Peng, T. hong; Lv, Y. cheng. Critical Roles of FTO-Mediated MRNA M6A Demethylation in Regulating Adipogenesis and Lipid Metabolism: Implications in Lipid Metabolic Disorders. Genes Dis. 2022, 9 (1), 51–61. 10.1016/j.gendis.2021.01.005.

(42) Borowski, T.; Bassan, A.; Siegbahn, P. E. M. Mechanism of Dioxygen Activation in 2-Oxoglutarate-Dependent Enzymes: A Hybrid DFT Study. Chem. - A Eur. J. 2004, 10 (4), 1031–1041. 10.1002/chem.200305306.

(43) Price, J. C.; Barr, E. W.; Hoffart, L. M.; Krebs, C.; Bollinger, J. M. Kinetic Dissection of the Catalytic Mechanism of Taurine:α-Ketoglutarate Dioxygenase (TauD) from Escherichia Coli. Biochemistry 2005, 44 (22), 8138–8147. 10.1021/bi050227c.

(44) Krebs, C.; Fujimori, D. G.; Walsh, C. T.; Bollinger, J. M. Non-Heme Fe (IV)– Oxo Intermediates. Acc. Chem. Res. 2007, 40 (7), 484–492.

(45) Bollinger Jr., J. M.; Chang, W.; Matthews, M. L.; Martinie, R. J.; Boal, A. K.; Krebs, C. Mechanisms of 2-Oxoglutarate-Dependent Oxygenases: The Hydroxylation Paradigm and Beyond. In 2-Oxoglutarate-Dependent Oxygenases; Schofield, C., Hausinger, R., Eds.; The Royal Society of Chemistry, 2015; p 0. 10.1039/9781782621959-00095.

(46) Martinez, S.; Hausinger, R. P. Catalytic Mechanisms of Fe(II)-and 2-Oxoglutarate-Dependent Oxygenases. J. Biol. Chem. 2015, 290 (34), 20702–20711. 10.1074/jbc.R115.648691.

(47) Price, J. C.; Barr, E. W.; Tirupati, B.; Bollinger, J. M.; Krebs, C. The First Direct Characterization of a High-Valent Iron Intermediate in the Reaction of an α-Ketoglutarate-Dependent Dioxygenase: A High-Spin Fe(IV) Complex in Taurine/α-Ketoglutarate Dioxygenase (TauD) from Escherichia Coli (Biochemistry (June 24, 2003). Biochemistry 2004, 43 (4), 1134. 10.1021/bi0330139.

(48) Vasta, J. D.; Raines, R. T. Human Collagen Prolyl 4-Hydroxylase Is Activated by Ligands for Its Iron Center. Biochemistry 2016, 55 (23), 3224–3233. 10.1021/acs.biochem.6b00251.

(49) Myllyla, R.; Kuutti-Savolainen, E. R.; Kivirikko, K. I. The Role of Ascorbate in the Prolyl Hydroxylase Reaction. Biochem. Biophys. Res. Commun. 1978, 83 (2), 441–448.

(50) De Jong, L.; Albracht, S. P. J.; Kemp, A. Prolyl 4-Hydroxylase Activity in Relation to the Oxidation State of Enzyme-Bound Iron. The Role of Ascorbate in Peptidyl Proline Hydroxylation. Biochim. Biophys. Acta (BBA)/Protein Struct. Mol. 1982, 704 (2), 326–332. 10.1016/0167-4838(82)90162-5.

(51) He, W.; Yin, X.; Xu, C.; Liu, X.; Huang, Y.; Yang, C.; Xu, Y.; Hu, L. Ascorbic Acid Reprograms Epigenome and Epitranscriptome by Reducing Fe(III) in the Catalytic Cycle of Dioxygenases. ACS Chem. Biol. 2024, 19 (1), 129–140. 10.1021/acschembio.3c00567.

(52) Smith-Díaz, C. C.; Das, A. B.; Jurkowski, T. P.; Hore, T. A.; Vissers, M. C. M. Exploring the Ascorbate Requirement of the 2-Oxoglutarate-Dependent Dioxygenases. J. Med. Chem. 2025. 10.1021/acs.jmedchem.4c02342.

(53) Myllyla, R.; Tuderman, L.; Kivirikko, K. I. Mechanism of the Prolyl Hydroxylase Reaction: 1. Role of Co-Substrates. Eur. J. Biochem. 1977, 80 (2), 349–357. 10.1111/j.1432-1033.1977.tb11889.x.

(54) Nelson, D. L.; Cox, M. M.; Hoskins, A. A. Lehninger Principles of Biochemistry, Eighth edi.; Macmillan Learning: New York, NY, 2021.

(55) Myllyharju, J. Prolyl 4-Hydroxylases, Key Enzymes in the Synthesis of Collagens and Regulation of the Response to Hypoxia, and Their Roles as Treatment Targets. Ann. Med. 2008, 40 (6), 402–417. 10.1080/07853890801986594.

(56) Yin, R.; Mao, S. Q.; Zhao, B.; Chong, Z.; Yang, Y.; Zhao, C.; Zhang, D.; Huang, H.; Gao, J.; Li, Z.; Jiao, Y.; Li, C.; Liu, S.; Wu, D.; Gu, W.; Yang, Y. G.; Xu, G. L.; Wang, H. Ascorbic Acid Enhances Tet-Mediated 5-Methylcytosine Oxidation and Promotes DNA Demethylation in Mammals. J. Am. Chem. Soc. 2013, 135 (28), 10396–10403. 10.1021/ja4028346.

(57) Welford, R. W. D.; Schlemminger, I.; McNeill, L. A.; Hewitson, K. S.; Schofield, C. J. The Selectivity and Inhibition of AlkB. J. Biol. Chem. 2003, 278 (12), 10157–10161. 10.1074/jbc.M211058200.

(58) Sun, H.; Li, K.; Zhang, X.; Liu, J.; Zhang, M.; Meng, H.; Yi, C. M6Am-Seq Reveals the Dynamic M6Am Methylation in the Human Transcriptome. Nat. Commun. 2021, 12 (1), 1–12. 10.1038/s41467-021-25105-5.

(59) Kuiper, C.; Molenaar, I. G. M.; Dachs, G. U.; Currie, M. J.; Sykes, P. H.; Vissers, M. C. M. Low Ascorbate Levels Are Associated with Increased Hypoxia-Inducible Factor-1 Activity and an Aggressive Tumor Phenotype in Endometrial Cancer. Cancer Res. 2010, 70 (14), 5749–5758. 10.1158/0008-5472.CAN-10-0263.

(60) Campbell, E. J.; Dachs, G. U.; Morrin, H. R.; Davey, V. C.; Robinson, B. A.; Vissers, M. C. M. Activation of the Hypoxia Pathway in Breast Cancer Tissue and Patient Survival Are Inversely Associated with Tumor Ascorbate Levels. BMC Cancer 2019, 19 (1), 1–13. 10.1186/s12885-019-5503-x.

(61) Jia, G.; Fu, Y.; Zhao, X.; Dai, Q.; Zheng, G.; Yang, Y.; Yi, C.; Lindahl, T.; Pan, T.; Yang, Y.; He, C. N6-Methyladenosine in Nuclear RNA Is a Major Substrate of the Obesity-Associated FTO. 2012, 7 (12), 885–887. 10.1038/nchembio.687.N.

(62) Zheng, G.; Dahl, J. A.; Niu, Y.; Fedorcsak, P.; Huang, C. M.; Li, C. J.; Vågbø, C. B.; Shi, Y.; Wang, W. L.; Song, S. H.; Lu, Z.; Bosmans, R. P. G.; Dai, Q.; Hao, Y. J.; Yang, X.; Zhao, W. M.; Tong, W. M.; Wang, X. J.; Bogdan, F.; Furu, K.; Fu, Y.; Jia, G.; Zhao, X.; Liu, J.; Krokan, H. E.; Klungland, A.; Yang, Y. G.; He, C. ALKBH5 Is a Mammalian RNA Demethylase That Impacts RNA Metabolism and Mouse Fertility. Mol. Cell 2013, 49 (1), 18–29. 10.1016/j.molcel.2012.10.015.

(63) Wei, J.; Liu, F.; Lu, Z.; Fei, Q.; Ai, Y.; He, P. C.; Shi, H.; Cui, X.; Su, R.; Klungland, A.; Jia, G.; Chen, J.; He, C. Differential m 6 A, m 6 A m, and m 1 A Demethylation Mediated by FTO in the Cell Nucleus and Cytoplasm. Mol. Cell 2018, 71 (6), 973-985.e5. 10.1016/j.molcel.2018.08.011.

(64) Zhang, X.; Wei, L. H.; Wang, Y.; Xiao, Y.; Liu, J.; Zhang, W.; Yan, N.; Amu, G.; Tang, X.; Zhang, L.; Jia, G. Structural Insights into FTO’s Catalytic Mechanism for the Demethylation of Multiple RNA Substrates. Proc. Natl. Acad. Sci. U. S. A. 2019, 116 (8), 2919–2924. 10.1073/pnas.1820574116.

(65) Kaur, S.; Tam, N. Y.; McDonough, M. A.; Schofield, C. J.; Aik, W. S. Mechanisms of Substrate Recognition and N6-Methyladenosine Demethylation Revealed by Crystal Structures of ALKBH5-RNA Complexes. Nucleic Acids Res. 2022, 50 (7), 4148–4160. 10.1093/nar/gkac195.

(66) Abramson, J.; Adler, J.; Dunger, J.; Evans, R.; Green, T.; Pritzel, A.; Ronneberger, O.; Willmore, L.; Ballard, A. J.; Bambrick, J.; Bodenstein, S. W.; Evans, D. A.; Hung, C. C.; O’Neill, M.; Reiman, D.; Tunyasuvunakool, K.; Wu, Z.; Žemgulytė, A.; Arvaniti, E.; Beattie, C.; Bertolli, O.; Bridgland, A.; Cherepanov, A.; Congreve, M.; Cowen-Rivers, A. I.; Cowie, A.; Figurnov, M.; Fuchs, F. B.; Gladman, H.; Jain, R.; Khan, Y. A.; Low, C. M. R.; Perlin, K.; Potapenko, A.; Savy, P.; Singh, S.; Stecula, A.; Thillaisundaram, A.; Tong, C.; Yakneen, S.; Zhong, E. D.; Zielinski, M.; Žídek, A.; Bapst, V.; Kohli, P.; Jaderberg, M.; Hassabis, D.; Jumper, J. M. Accurate Structure Prediction of Biomolecular Interactions with AlphaFold 3. Nature 2024, 630 (8016), 493–500. 10.1038/s41586-024-07487-w.

(67) Alves, J.; Vidugiris, G.; Goueli, S. A.; Zegzouti, H. Bioluminescent High-Throughput Succinate Detection Method for Monitoring the Activity of JMJC Histone Demethylases and Fe(II)/2-Oxoglutarate-Dependent Dioxygenases. SLAS Discov. 2018, 23 (3), 242–254. 10.1177/2472555217745657.

(68) Šimelis, K.; Belle, R.; Kawamura, A. Unravelling 2-Oxoglutarate Turnover and Substrate Oxidation Dynamics in 5-Methylcytosine-Oxidising TET Enzymes. Commun. Chem. 2024, 7 (1), 1–9. 10.1038/s42004-024-01382-1.

(69) Komadel, P.; Stucki, J. W. Quantitative Assay of Minerals for Fe2 + and Fe3 + Using 1,10-Phenanthroline: Iii. a Rapid Photochemical Method. Clays Clay Miner. 1988, 36 (4), 379–381. 10.1346/CCMN.1988.0360415.

(70) Purslow, J. A.; Nguyen, T. T.; Egner, T. K.; Dotas, R. R.; Khatiwada, B.; Venditti, V. Active Site Breathing of Human Alkbh5 Revealed by Solution NMR and Accelerated Molecular Dynamics. Biophys. J. 2018, 115 (10), 1895–1905. 10.1016/j.bpj.2018.10.004.

(71) Burns, D.; Khatiwada, B.; Singh, A.; Purslow, J. A.; Potoyan, D. A.; Venditti, V. An α-Ketoglutarate Conformational Switch Controls Iron Accessibility, Activation, and Substrate Selection of the Human FTO Protein. Proc. Natl. Acad. Sci. 2024, 121 (25), e2404457121. 10.1073/pnas.2404457121.

(72) Wang, T.; Hong, T.; Huang, Y.; Su, H.; Wu, F.; Chen, Y.; Wei, L.; Huang, W.; Hua, X.; Xia, Y.; Xu, J.; Gan, J.; Yuan, B.; Feng, Y.; Zhang, X.; Yang, C.-G.; Zhou, X. Fluorescein Derivatives as Bifunctional Molecules for the Simultaneous Inhibiting and Labeling of FTO Protein. J. Am. Chem. Soc. 2015, 137 (43), 13736–13739. 10.1021/jacs.5b06690.

(73) Li, W.; Zhang, T.; Sun, M.; Shi, Y.; Zhang, X. J.; Xu, G. L.; Ding, J. Molecular Mechanism for Vitamin C-Derived C5-Glyceryl-Methylcytosine DNA Modification Catalyzed by Algal TET Homologue CMD1. Nat. Commun. 2021, 12 (1), 1–13. 10.1038/s41467-021-21061-2.

(74) Sato, N.; Uragami, Y.; Nishizaki, T.; Takahashi, Y.; Sazaki, G.; Sugimoto, K.; Nonaka, T.; Masai, E.; Fukuda, M.; Senda, T. Crystal Structures of the Reaction Intermediate and Its Homologue of an Extradiol-Cleaving Catecholic Dioxygenase. J. Mol. Biol. 2002, 321 (4), 621–636. 10.1016/S0022-2836(02)00673-3.

(75) Imparato, C.; D’Errico, G.; Macyk, W.; Kobielusz, M.; Vitiello, G.; Aronne, A. Interfacial Charge Transfer Complexes in TiO2-Enediol Hybrids Synthesized by Sol-Gel. Langmuir 2022, 38 (5), 1821–1832. 10.1021/acs.langmuir.1c02939.

(76) Ang, A.; Pullar, J. M.; Currie, M. J.; Vissers, M. C. M. Vitamin C and Immune Cell Function in Inflammation and Cancer. Biochem. Soc. Trans. 2018, 46 (5), 1147–1159. 10.1042/BST20180169.

(77) Warminski, M.; Trepkowska, E.; Smietanski, M.; Sikorski, P. J.; Baranowski, M. R.; Bednarczyk, M.; Kedzierska, H.; Majewski, B.; Mamot, A.; Papiernik, D.; Popielec, A.; Serwa, R. A.; Shimanski, B. A.; Sklepkiewicz, P.; Sklucka, M.; Sokolowska, O.; Spiewla, T.; Toczydlowska-Socha, D.; Warminska, Z.; Wolosewicz, K.; Zuberek, J.; Mugridge, J. S.; Nowis, D.; Golab, J.; Jemielity, J.; Kowalska, J. Trinucleotide MRNA Cap Analogue N6-Benzylated at the Site of Posttranscriptional M6Am Mark Facilitates MRNA Purification and Confers Superior Translational Properties In Vitro and In Vivo. J. Am. Chem. Soc. 2024, 146 (12), 8149–8163. 10.1021/jacs.3c12629.

(78) Smart, O.S., Sharff A., Holstein, J., Womack, T.O., Flensburg, C., Keller, P., Paciorek, W., Vonrhein, C. and Bricogne G. (2021) Grade2 version 1.7.0. Cambridge, United Kingdom: Global Phasing Ltd.

