## Supplementary Materials for "How ascorbate tunes the activity and substrate selectivity of Fe(II)-dependent dioxygenase superfamily enzymes"

**This PDF file includes:**

Figures S1 to S4
Tables S1 to S3

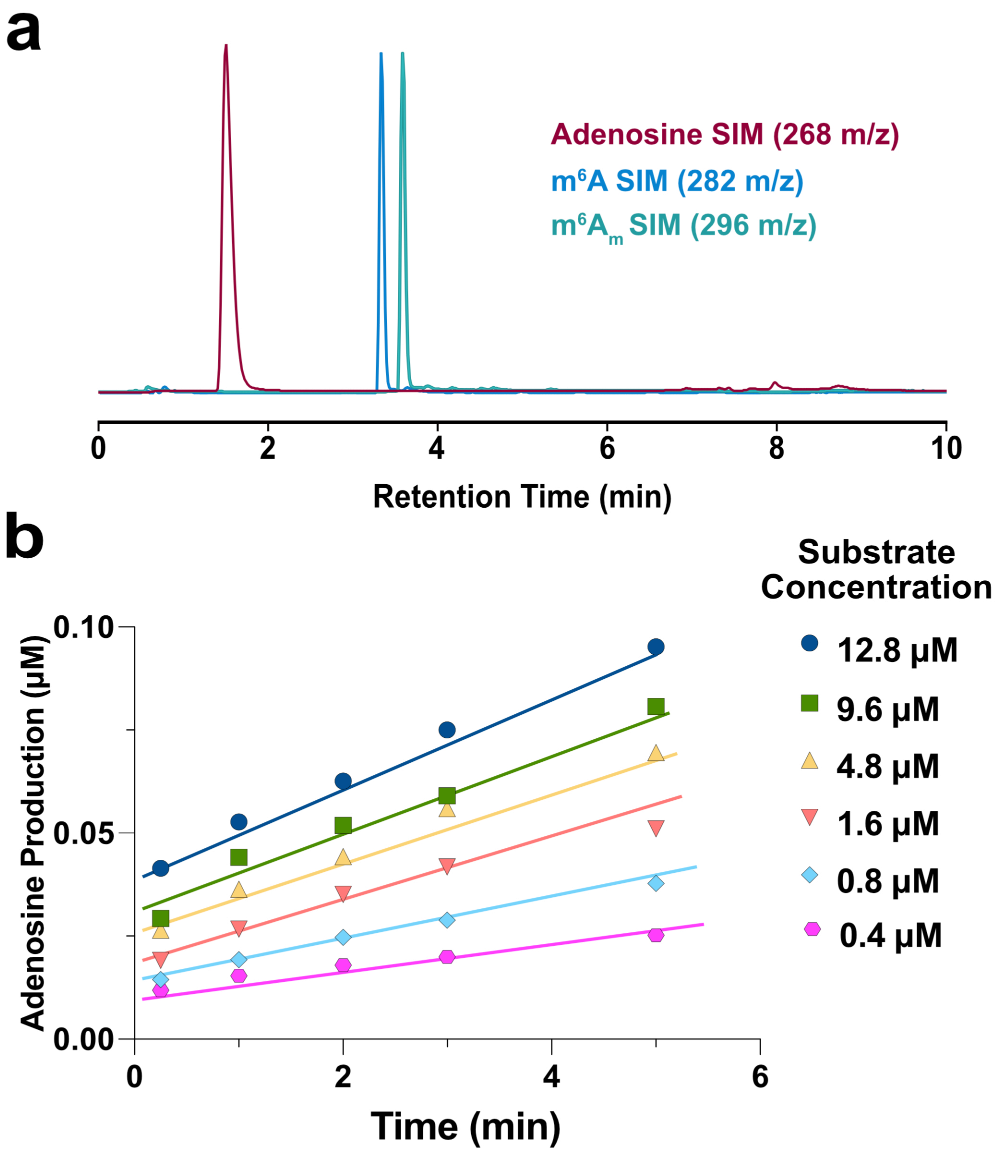

**Supplementary Figure 1. Representative m$^6$A-RNA demethylation kinetic analysis.** (**a**) UHPLC-MS traces showing normalized SIMs of adenosine, $A_m$, and m$^6A_m$ commercial nucleoside standards. Distinct nucleoside peaks from quenched reaction samples are integrated and compared to a dilution series of nucleoside standards to quantify nucleoside levels. (**b**) Representative product adenosine concentration versus time plots used to obtain initial rates for kinetic analyses (data shown for AlkBH5 with m$^6$A 9-mer substrate). Initial rates were calculated from the slope of the linear trendline in 5-minute time courses.

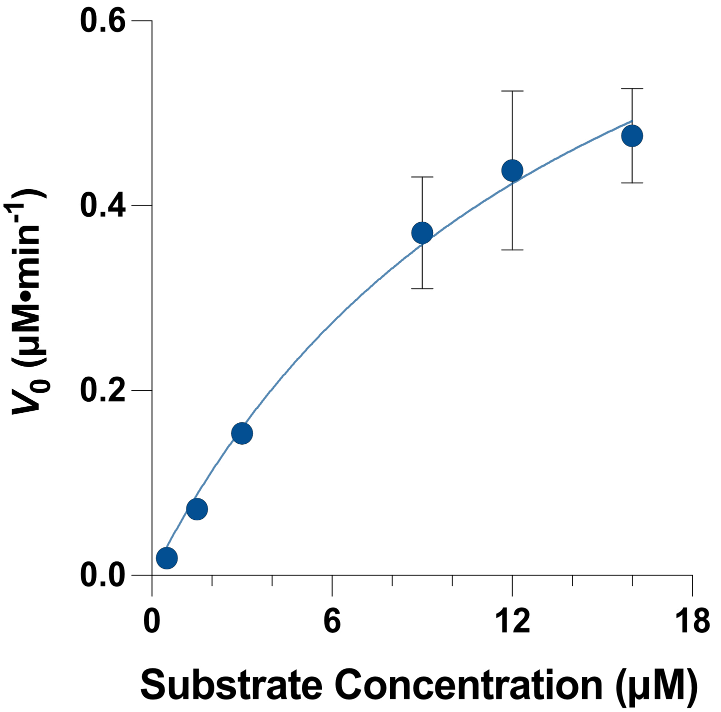

**Supplementary Figure 2. Michaelis-Menten plot for FTO demethylation of m$^6$A$_m$ cap RNA substrate.** Initial rates of *in vitro* FTO-mediated demethylation reactions are plotted for different m$^6$A$_m$-cap RNA substrate concentrations and fit to the Michaelis-Menten equation to determine kinetic parameters. FTO (200 nM) was incubated with Fe(II), 2-OG, and varied concentrations of m$^6$A$_m$-cap RNA substrate. Error bars represent SEM from three replicates. Kinetic parameters are listed in **Supplementary Table 1a**.

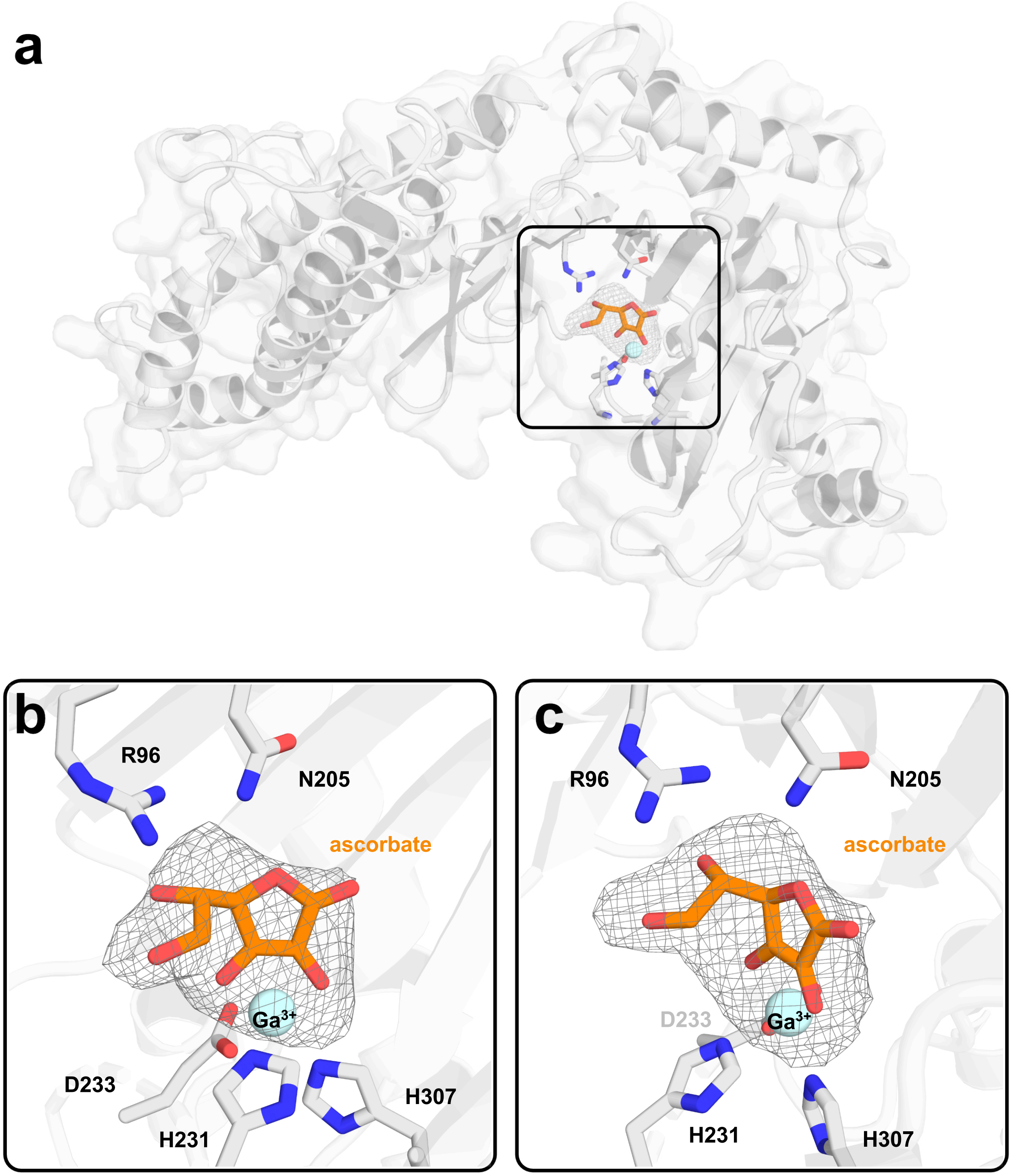

**Supplementary Figure 3. Structure of FTO with ascorbate. (a)** Full FTO structure with boxed active site area showing ascorbate $F_O$-$F_C$ omit map and key FTO active site residues as sticks. **(b, c)** Zoom-in of boxed active site area in (a) showing two different views of the ascorbate $F_O$-$F_C$ omit map and modeled ascorbate (orange sticks) bound to $Ga^{3+}$ metal (blue sphere). $F_O$-$F_C$ omit map is shown at 3.0σ.

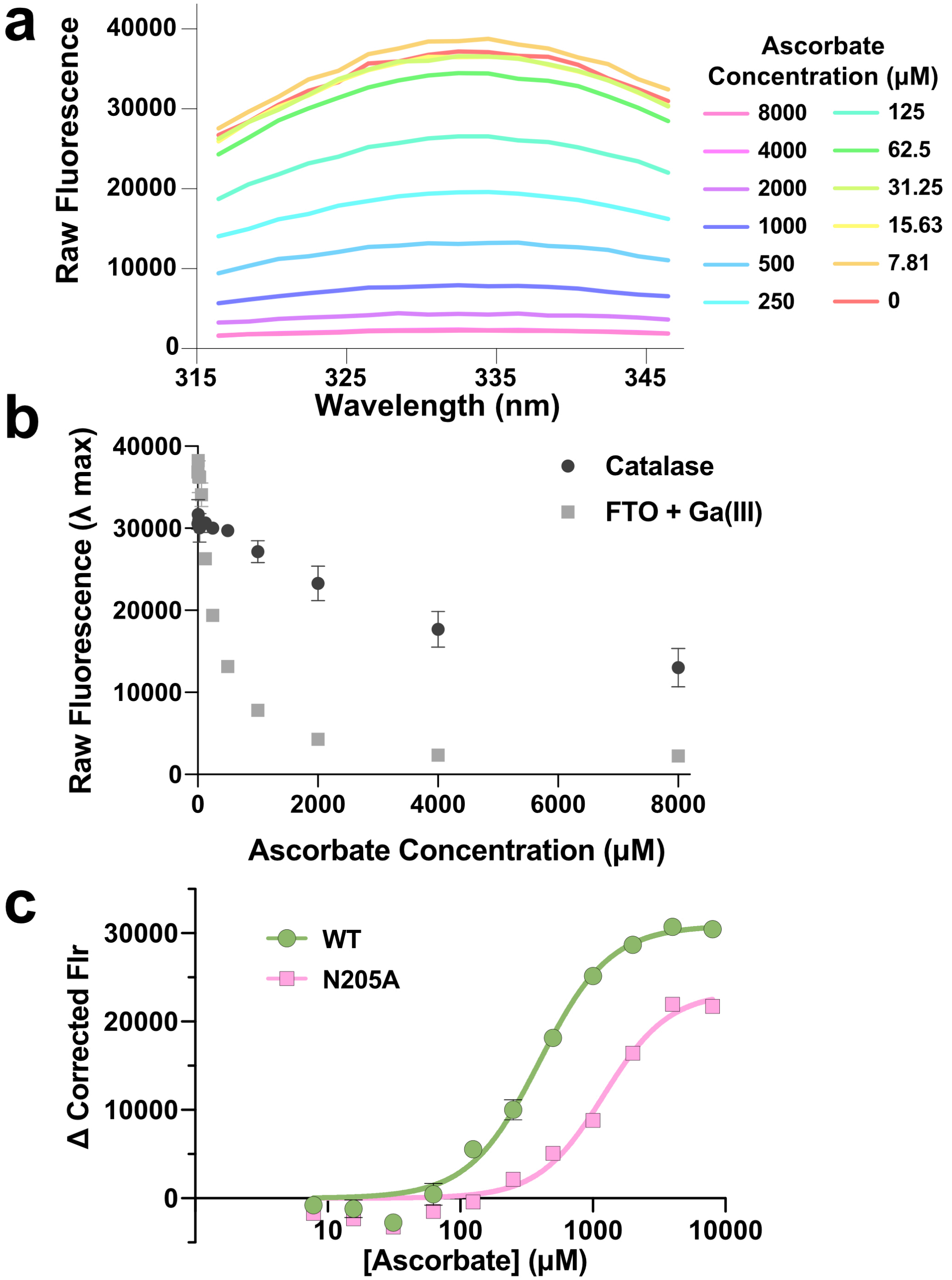

**Supplementary Figure 4. Intrinsic tryptophan fluorescence binding data. (a)** Representative tryptophan fluorescence spectra of WT FTO + Ga(III) solutions with varying ascorbate concentration; only the Trp fluorescence peak maximum (~ 315 – 345 nm) is shown. Increasing ascorbate concentrations quench the tryptophan fluorescence peak. **(b)** Raw Trp fluorescence data collected from ascorbate titrations with WT FTO + Ga(III) (gray) or catalase control (black). The catalase control was used to correct for non-specific binding and inner filter effects, as outlined in the Methods section. **(c)** Corrected fluorescence binding curves for titration of ascorbate into WT FTO + Mn(II) (green) or FTO N205A + Mn(II) (pink).

**Supplementary Table 1. Kinetic parameters for FTO- and AlkBH5-mediated demethylation reactions.** (**a**) Kinetic parameters calculated from initial rates of FTO reactions at 37 °C with $m^6A$ linear 9-mer, $m^6A$ 25-mer stem-loop, and $m^6A_m$-cap RNA substrates *in vitro,* all with 2 mM ascorbate. Kinetic parameters for FTO demethylation reactions in the absence of ascorbate were not determined due to low activity with $m^6A$ RNA substrates. (**b**) Kinetic parameters calculated from initial rates of AlkBH5 reactions at 30 °C with $m^6A$ linear 9-mer and $m^6A$ 25-mer stem-loop RNA substrates *in vitro*, with either 2 mM ascorbate (+Asc.) or 0 mM ascorbate (-Asc.). A lower temperature was used for AlkBH5 reactions in order to be able to capture initial rates by manual quenching. We note that although the different reaction temperatures needed to capture AlkBH5 versus FTO initial rates makes cross comparison of their kinetic parameters difficult, all the comparative conclusions drawn from these data are exclusively comparing kinetic parameters of different substrates for FTO, or different substrates for AlkBH5, not comparing kinetic parameters for FTO versus AlkBH5.

**a**

| FTO: Substrate | $K_M$ (μM) | $k_{cat}$ ($min^{-1}$) | $k_{cat}/K_M$ ($μM^{-1} • min^{-1}$) |
|---|---|---|---|
| $m^6A$ Linear 9-mer (+ Asc.) | 23.87 +/- 5.11 | 0.02 +/- 0.002 | 0.0008 ± 0.0002 |
| $m^6A$ 25-mer Stem-Loop (+ Asc.) | 4.21 +/- 1.71 | 0.32 +/- 0.04 | 0.076 ± 0.032 |
| $m^6A_m$ Cap (+ Asc.) | 14.81 +/- 8.88 | 4.73 +/- 1.6 | 0.319 ± 0.220 |

**b**

| AlkBH5: Substrate | $K_M$ (μM) | $k_{cat}$ ($min^{-1}$) | $k_{cat}/K_M$ ($μM^{-1} • min^{-1}$) |
|---|---|---|---|
| $m^6A$ Linear 9-mer (+ Asc.) | 1.02 +/- 0.22 | 0.06 +/- 0.003 | 0.059 ± 0.013 |
| $m^6A$ Linear 9-mer (- Asc.) | 0.41 +/- 0.08 | 0.05 +/- 0.002 | 0.122 ± 0.024 |
| $m^6A$ 25-mer Stem-Loop (+ Asc.) | 2.73 +/- 0.45 | 0.07 +/- 0.004 | 0.026 ± 0.005 |
| $m^6A$ 25-mer Stem-Loop (- Asc.) | 7.12 +/- 2.06 | 0.07 +/- 0.009 | 0.01 ± 0.003 |

**Supplementary Table 2.** Data and refinement statistics for crystal structure of FTO in complex with Ga(III) and ascorbate (PDB 9OHS).

| | **Crystal structure of human FTO in complex with Ga(III) and ascorbate (PDB 9OHS)** |
|---|---|
| **Data collection** | |
| Space group | H3 |
| Cell dimensions | |
| $a$, $b$, $c$ (Å) | 143.872, 143.872, 85.241 |
| □□□□□□□□□□□□□ (□) | 90.00, 90.00, 120.00 |
| Resolution (Å) | 41.22 – 3.07 (3.18 – 3.07)[a] |
| $R_{\mathrm{merge}}$ | 0.1285 (1.629) |
| $I$ / σ$I$ | 14.73 (1.31) |
| $CC_{1/2}$ | 0.999 (0.283) |
| Completeness (%) | 99.91 (99.84) |
| Multiplicity | 10.7 (11.1) |
| **Refinement** | |
| Resolution (Å) | 41.22 – 3.07 (3.18 – 3.07) |
| No. reflections | 12,274 (1,235) |
| $R$ / $R_{\mathrm{free}}$ | 0.1929 / 0.2529 |
| No. non-H atoms | |
| Protein | 3475 |
| Ligand | 13 |
| $B$-factors | |
| Protein | 121.35 |
| Ligand | 134.34 |
| R.m.s. deviations | |
| Bond lengths (Å) | 0.011 |
| Bond angles (□) | 1.25 |
| Ramachandran plot statistics | |
| No. favored | 385 (90.2 %) |
| No. allowed | 42 (9.8 %) |
| No. outliers | 0 (0.0 %) |

Data set was collected from a single crystal. [a]Values in parentheses are for highest-resolution shell.

**Supplementary Table 3. Ascorbate binding parameters determined from intrinsic tryptophan fluorescence titrations.** $K_D$ values for ascorbate binding to FTO + metal were obtained by fitting corrected fluorescence data to a standard Hill binding model in GraphPad Prism as described in the Methods section.

| FTO construct with metal | $K_D$ (μM) ± (SEM) |
|---|---|
| WT + Ga(III) | 201.2 ± 10.0 |
| N205A + Ga(III) | 830.1 ± 79.1 |
| WT + Mn(II) | 394.5 ± 27.8 |
| N205A + Mn(II) | 1207.9 ± 136.2 |
